# Structural details of helix-mediated TDP-43 C-terminal domain multimerization

**DOI:** 10.1101/2024.07.05.602258

**Authors:** Azamat Rizuan, Jayakrishna Shenoy, Priyesh Mohanty, Patricia M. S. dos Passos, José F. Mercado Ortiz, Leanna Bai, Renjith Viswanathan, Szu-Huan Wang, Victoria Johnson, Lohany D. Mamede, Yuna M. Ayala, Rodolfo Ghirlando, Jeetain Mittal, Nicolas L. Fawzi

## Abstract

The primarily disordered C-terminal domain (CTD) of TAR DNA binding protein-43 (TDP-43), a key nuclear protein in RNA metabolism, forms neuronal inclusions in several neurodegenerative diseases. A conserved region (CR, spanning residues 319-341) in CTD forms transient helix-helix contacts important for its higher-order oligomerization and function that are disrupted by ALS-associated mutations. However, the structural details of CR assembly and the explanation for several ALS-associated variants’ impact on phase separation and function remain unclear due to challenges in analyzing the dynamic association of TDP-43 CTD using traditional structural biology approaches. By employing an integrative approach, combining biophysical experiments, biochemical assays, AlphaFold2-Multimer (AF2-Multimer), and atomistic simulations, we generated structural models of helical oligomerization of TDP-43 CR. Using NMR, we first established that the native state of TDP-43 CR under physiological conditions is α-helical. Next, alanine scanning mutagenesis revealed that while hydrophobic residues in the CR are important for CR assembly, phase separation and TDP-43 nuclear retention function, polar residues down regulate these processes. Finally, pairing AF2-Multimer modeling with AAMD simulations indicated that dynamic, oligomeric assemblies of TDP-43 that are stabilized by a methionine-rich core with specific contributions from a tryptophan/leucine pair. In conclusion, our results advance the structural understanding of the mechanisms driving TDP-43 function and provide a window into the initial stages of its conversion into pathogenic aggregates.

## Introduction

Transactive response DNA-binding protein 43 (TDP-43) is a nucleo-cytoplasmic shuttling RNA-binding protein (RBP), with predominant nuclear localization, engaging in diverse cellular functions across both the nucleus and cytoplasm, including RNA transcription, splicing, and transport^1, 2^. Many TDP-43 functions are linked to the formation of dynamic, intracellular membraneless organelles via phase separation^3–6^. Furthermore, TDP-43 is identified as a major component of ubiquitinated neuronal inclusions in the cytoplasm, a pathological hallmark of amyotrophic lateral sclerosis (ALS), frontotemporal dementia (FTD), and limbic-predominant age-related TDP-43 encephalopathy (LATE), a condition with pathology and symptoms overlapping with Alzheimer’s disease^7–11^. The protein consists of a folded N-terminal domain (NTD) responsible for dynamic homo-oligomerization^12, 13^, two RNA recognition motifs (RRMs) that together bind GU-rich RNA sequences^14, 15^, and an intrinsically disordered C-terminal domain (CTD) characterized by a low-complexity (LC) sequence^16, 17^. The aggregation-prone CTD is a hotspot for ∼90% of all ALS-associated mutations^18, 19^, and proteolytically cleaved C-terminal fragments are enriched in pathological inclusions^7^, underscoring its importance in TDP-43 dysfunction. In biochemical assays, the CTD is sufficient for phase separation in the presence of physiological salt concentrations or RNA^16^. Additionally, TDP-43 CTD condensation is promoted by the hydrophobic conserved region (CR, spanning residues 319-341), working in concert with the adjacent IDRs^5, 20^.

The CR in CTD is uniquely well-conserved throughout vertebrates compared to the flanking IDRs^16, 21^ (**Fig. 1A**). Previous work from our laboratories^16^ and others^22^ show that the CR adopts a transient α-helical structure under physiological conditions. Furthermore, the CR mediates intermolecular helix-helix contacts that are essential for liquid-like phase separation of CTD *in vitro*^16^, phase separation of TDP-43 reporter constructs in cells^21^, splicing^5^, 3’ polyadenylation function^3^, TDP-43 RNA-level autoregulation^3^, and the cytosolic TDP-43 aggregation under oxidative stress^23^. The disruption of these interactions by ALS-associated mutations or by engineered helix-disrupting variants within the CR can perturb TDP-43 phase separation^5, 16^ and can lead to aggregation via structural conversion of the CR into β-sheet aggregates in disease^9, 10, 23, 24^. Conversely, helical self-assembly can be enhanced by designed CR variants, enabling control over the function and material properties of phase-separated droplets^5^. Using NMR experiments combined with computer simulations, we previously observed that α-helicity within the CR is enhanced during helix-helix dimerization, leading to significant structural changes^5^.

**Figure 1.**
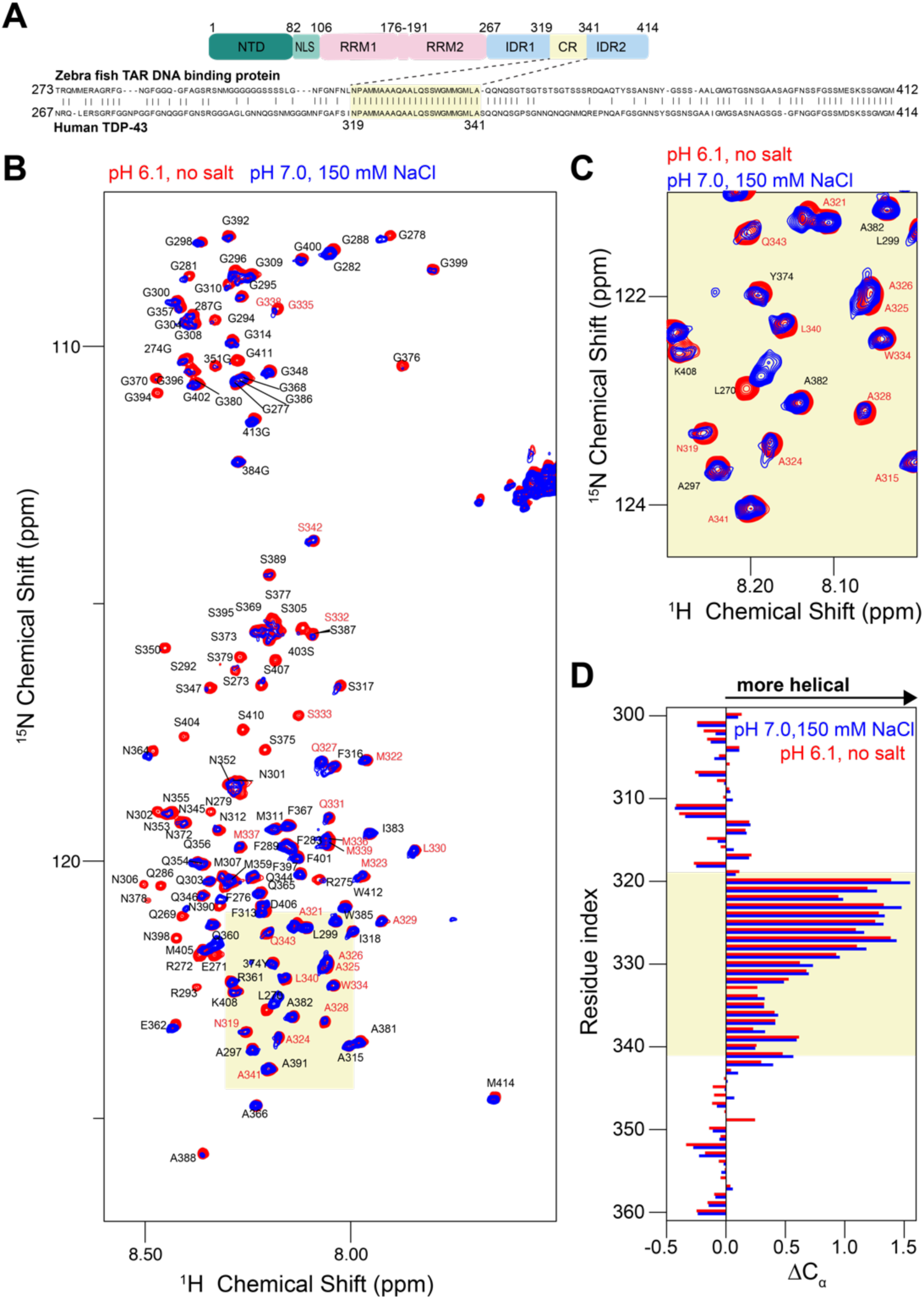
The conserved region (CR) of TDP-43 CTD adopts α-helical structure at native buffer conditions. **A.** Domain composition of TDP-43, with CR residues highlighted. Pairwise alignment of the CTD of human TDP-43 (UniProt id: Q13148) and zebrafish TAR DNA binding protein (UniProt id: Q802C7) shows CR residues are evolutionarily conserved in vertebrates. Identical residues are indicated with vertical bars. **B.** A comparison of the ^1^H- ^15^N HSQC of WT CTD at NMR conditions (20 mM MES, pH 6.1, red) and neutral pH, with physiologically relevant monovalent salt concentration (20 mM HEPES, 150 mM NaCl buffer pH 7, blue) shows very similar spectral fingerprint, especially in the CR region (red text). **C**. An excerpt of the ^1^H-^15^N HSQC shows resonances from the CR are nearly identical at both conditions. **D.** Secondary C_α_ chemical shifts comparing the observed chemical shift and the value for a random coil reference, ΔδC_α_, at pH 6.1 without salt and pH 7.0 with 150 mM NaCl, showing that the helicity is observed in both conditions. Positive ΔCα values are indicative of α-helical secondary structure.

Most importantly, the basic stoichiometry of TDP-43 CR interactions has proved elusive. Although definitive evidence has yet to emerge, the presence of helix-mediated higher-order oligomers larger than dimers has been postulated based on several lines of evidence including NMR diffusion data as a function of CTD concentration^5^ and elevated NMR relaxation values extracted from chemical exchange experiments^16^. However, the molecular size of these self-assembled species and their atomic structural details remain unresolved due to rapid aggregation of CTD at higher concentrations.

Here, we employed an integrative modeling strategy based on extensive experimental characterization using NMR spectroscopy and sedimentation velocity analytical ultracentrifugation (SV-AUC), coupled with state-of-the-art computational methods such as AF2-Multimer^25, 26^ and all-atom molecular dynamics (AAMD) simulations, to probe the atomic structural details of TDP-43 CR helix-mediated assembly. Importantly, we establish atomic-level structural models of TDP-43 CR multimeric assemblies observed in SV-AUC experiments which provides critical information required to understand the early steps in TDP-43 CTD assembly. These structural insights constitute a significant advancement in our comprehension of the molecular pathways underlying TDP-43 phase separation and function, thereby opening avenues for studying the emergence of its pathogenicity.

## Results and Discussion

### The native state of TDP-43 CR at physiological conditions is α-helical

Previously, we provided direct structural evidence for the presence of α-helical structure within CR of TDP-43 and its stabilization via helix-helix assembly using solution NMR spectroscopy^5, 16^. The experiments were performed at a slightly acidic pH (6.1) and without salt to attain optimal NMR spectra and prevent phase separation. However, recent work using hydrogen-bond disrupting variants suggested (without direct structural evidence) that the CR of TDP-43 adopts a β-sheet conformation at neutral pH and physiological salt and that the α-helical structure previously observed by us and others was an artifact of non-physiological conditions. To resolve this important issue, we conducted solution NMR spectroscopic measurements at both our previous conditions (slightly acidic pH 6.1 with no salt) and at physiological conditions (neutral pH 7.0 with 150 mM NaCl) and low protein concentration to prevent phase separation.

The largest differences in the spectra come from the loss of several resonances for serine and glycine residues in the IDR regions, (**Fig. 1B**), due to rapid exchange of the amide hydrogen for these residue types at higher pH^27^. Focusing on the overlay of ^1^H-^15^N HSQC spectra for residues within the CR (**Fig. 1C**), we observed only minimal changes in chemical shifts, suggesting no significant change in the overall structure of this region. To further probe structural changes, we compared the differences between experimental C_α_ chemical shifts and the random coil shifts, ΔδC_α_ (**Fig. 1D**)^28, 29^, and found highly similar ΔδC_α_ values in both cases. Positive ΔδC_α_ values within the CR reveal its helical character at neutral pH and physiological salt conditions; in fact, helicity is slightly enhanced at these conditions. Furthermore, our previous computer simulations, performed under physiological salt and pH conditions using a modern state-of-the-art force field for intrinsically disordered proteins^30^, also confirmed the presence of a transient helix spanning residues 321-330 in the monomeric CTD ensemble and its stabilization by intermolecular interactions^5, 16, 20^. These observations are also consistent with solid-state NMR experiments on *in vitro* aged liquid droplets^31^ and fibrillar aggregates^32, 33^, which showed the presence of helical structures in TDP-43 CR even after phase transition to fibrils^33^. Taken together, our direct structural measurements using NMR unequivocally show the presence of native α-helical structure within TDP-43 CTD CR at physiological conditions.

### Alanine-scanning mutagenesis within TDP-43 CR reveal key residues in phase separation

TDP-43 CR promotes the formation of phase-separated CTD droplets^16^. Disease and designed mutations that reduce helical stability or disrupt intermolecular helix-helix interactions decrease CTD phase separation, while variants enhancing helicity promote it^5^. By simultaneous substitution of all five CR methiniones to alanine, we recently found that methionine residues within CR are crucial for intermolecular helix-helix contacts mediating CTD phase separation^20^. To comprehensively investigate the role of each CR position, we created TDP-43 CTD variants substituting each non-alanine position with alanine to preserve helicity while potentially disrupting CR:CR interactions^20^. We measured the saturation concentration (*c*_sat_), the protein concentration threshold for phase separation, using droplet sedimentation assays for all variants. Microscopy was used to evaluate whether droplets remained dynamic and liquid-like or transitioned to static aggregates.

In the absence of salt, WT CTD did not undergo phase separation but formed liquid-like droplets with increasing salt concentrations, having a *c*_sat_ ∼15 μM at 150 mM NaCl (**Fig. 2A,B, Fig. S1**). Highlighting the essential role of a single tryptophan in promoting CTD condensation, W334A showed the most profound effect on phase separation, resulting in no droplets at the studied conditions (**Fig. 2B**). Similarly, *c*_sat_ was increased (decreased phase separation) three-fold by F316A which lies directly adjacent to the helical CR, whereas the effect of F313A in the region preceding the helical CR was less, showing *c*_sat_ twice as high as that of WT (**Fig. 2A**).

**Figure 2.**
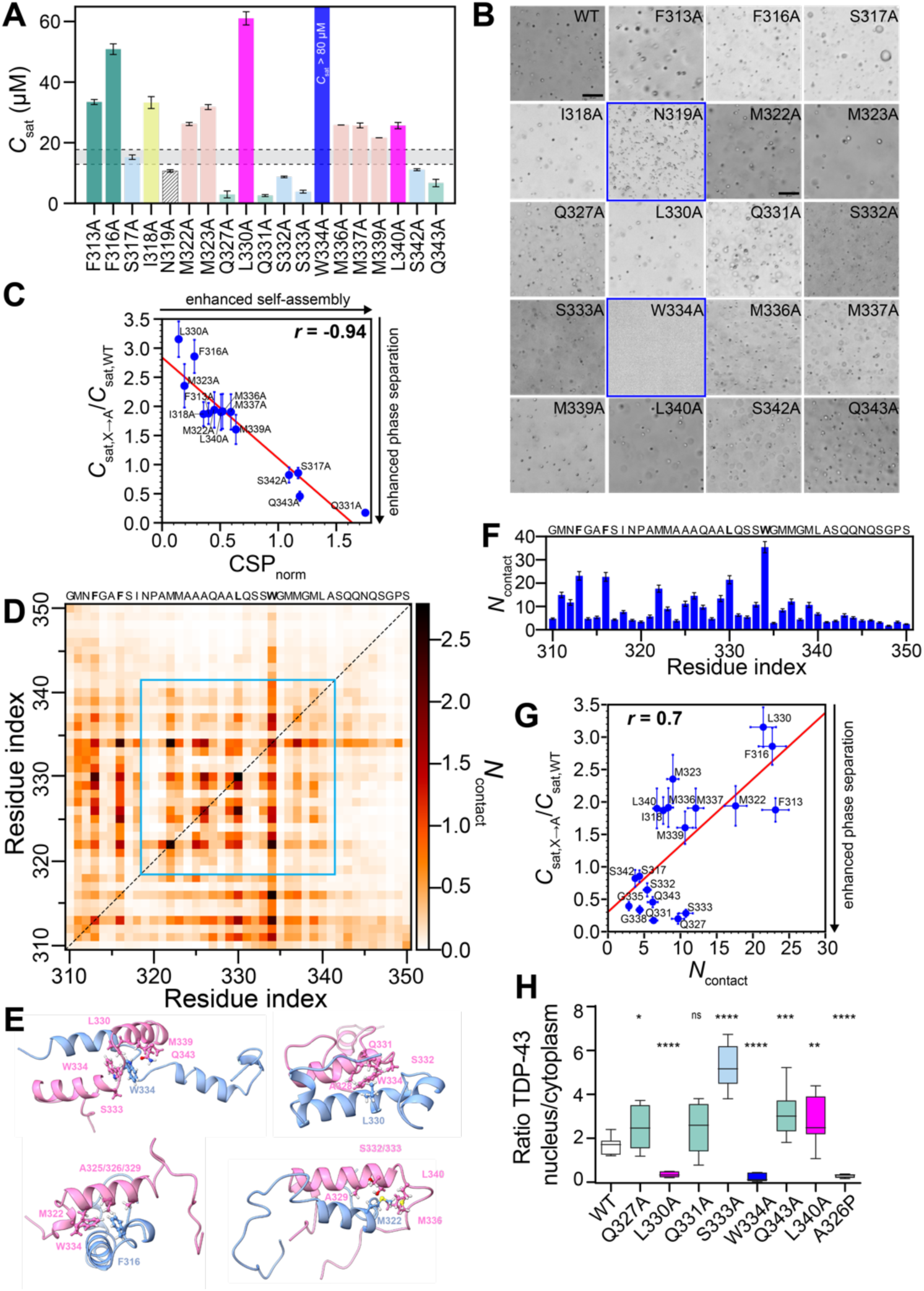
Site-specific alanine substitutions within the TDP-43 CR reveal key residues in TDP-43 phase separation, helix assembly, and function. **A.** Saturation concentration (*c*_sat_) for WT CTD (represented by a grey box with dashed lines covering the mean ± standard deviation from independent experiments shown in Fig. S1) and its single alanine substitution variants at non-alanine positions within CR and its adjacent residues are determined by measuring the protein remaining in supernatant after phase separation at 150 mM NaCl. **B**. DIC micrographs of CTD variants at 80 μM in 150 mM salt buffer. Scale bar, 20 μm. All variants form round liquid-like droplets except N319A that forms morphologically irregular assemblies and W334A that does not form droplets at these conditions, highlighted with a blue box. **C.** ^15^N chemical shift perturbations (CSP) (Δδ^15^N) and *c*_sat_ for the single alanine substitution variants exhibit a strong correlation (Pearson correlation coefficient, r = −0.94). Both axes is normalized by WT CTD values. **D**. Pairwise intermolecular contact map as a function of residue index from two-chain AAMD simulations of TDP-43_310–350_ at 150 mM NaCl. The contact maps are averaged over 18 independent trajectories (total runtime ∼100 μs). Contacts within CR are highlighted with a cyan box. **E.** Representative snapshots from two-chain AAMD simulations of TDP-43_310–350_ at 150 mM NaCl show dynamically interconverting structures with heterogeneous helix-helix contacts, involving a multitude of residues. **F.** Total number of contacts (N_contact_) for each residue position computed by summation of all pairwise contacts from the two dimensional pairwise intermolecular contact map from D. **G.** Per-residue intermolecular contacts between TDP-43_310–350_ pairs in two-chain simulations show a good correlation (Pearson correlation coefficient, *r* = 0.7) with the *c*_sat_ from single alanine substitution variants at non-alanine positions within CR and its adjacent residues (including G335A and G338A *C*_sat_ values from our previous work^5^), normalized by *c*_sat_ of WT CTD. **H.** TDP-43 nuclear to cytoplasmic localization ratio (N/C) of WT TDP-43 and its single alanine substitution variants within CR in HEK293^HA-TDP-43^ quantified from immunoblotting. N/C ratio of helix-disrupting mutant, A326P is included as a control. \*\*\**p* < 0.001, \**p* < 0.05, ns (not significant) p > 0.05, one-way ANOVA with Tukey’s test.

Expanding from our previous results for the 5M→A variant within CR, which collectively impaired the CTD phase separation, each single methionine substitution within CR (M322A, M333A, M336A, M337A, M339A) also reduce phase separation with approximately two-fold higher *c*_sat_ than WT (**Fig. 2A**). Similarly, I318A and L340A result in *c*_sat_ values similar to single methionine to alanine substitutions, suggesting that bulky aliphatic residues contribute similarly to phase separation. Somewhat surprisingly, the substitution of L330A emerges as a major outlier compared to other aliphatic residues, resulting in significant destabilization of the condensate (*c*_sat_ ∼ 60 μM). This effect is much higher than L340A near the C-terminal end of the CR, suggesting a special role of L330 in self-assembly.

Next, we consider the polar residues which are just as conserved as the hydrophobic residues in the CR (**Fig. 2A**). Alanine mutations at most serine positions (S317A, S332A, S342A) exhibit small enhancements in phase separation compared to WT CTD, except for S333A which shows a marked enhancement (∼four-fold) in *c*_sat_. Glutamine to alanine mutations (Q327A, Q331A, Q343A) display the greatest enhancement of phase separation with *c*_sat_ ∼5 μM, approximately three-fold lower than WT *c*_sat_. Microscopy of these variants after phase separation confirm no morphological differences compared to WT droplets and no evidence of a conversion to solid-like aggregates, except for N319A, which results in morphologically irregularly shaped assemblies. Similar to how glycine-to-alanine substitutions (G335A, G338A) enhance CTD self-association and phase separation by stabilizing the helicity of the monomeric CR^5^, the enhancement in phase separation for polar-to-alanine substitutions reinforce a critical role of the CR helix. Taken together, *in vitro* phase separation experiments highlight the importance of multiple residues within CR for phase separation, with phase separation promotion in particular by L330, W334, and methionines while polar resiudes either down modulate phase separation or prevent aggregation.

### TDP-43 CR helix-mediated interactions are linked to CTD phase separation

To probe the relationship between helix-mediated interactions and phase separation, we evaluated the impact of alanine mutations on the helix-helix contacts by monitoring NMR chemical shift perturbations (CSPs) as a function of protein concentration under conditions of pH 6.1 and without salt, where TDP-43 CTD does not readily phase separate. The relative magnitude of helix-mediated self-assembly for each variant was quantified by the ^15^N CSP (Δδ^15^N) for the A328 resonance at 90 μM (where intermolecular interactions are formed) and 20 μM (monomeric reference state), normalized to WT CTD values (CSP_norm_). Q327A, S332A, and S333A were excluded due to broadening of CR signals beyond detection in HSQC experiments, consistent with their enhanced self-assembly. W334A and N319A were also excluded as W334A does not phase separate, and N319A results in irregularly shaped aggregates, making the apparent *c*_sat_ unreliable. Remarkably, we observed a strong correlation between CSP_norm_ and *c*_sat_ for alanine mutants, with a Pearson correlation coefficient (*r*) of −0.94 (**Fig. 2C**), i.e., higher CSP_norm_ values were observed for alanine variants enhancing phase separation and lower values for those reducing phase separation. Together, our data show that helix-mediated interactions are linked to the phase separation of TDP-43 CTD.

To understand the residue-level molecular interactions driving intermolecular helix-mediated assembly, we performed dimer structure predictions for a TDP-43 subregion containing the CR (residues 310-350) using AF2-Multimer (**Fig. S2A**). The predicted dimer states showed high structural heterogeneity and low model confidence, suggesting that CR may not form a stable, stereospecific dimer and instead undergo dynamic interconversion with respect to the monomeric state. (**Fig. S2B**). To test this, we conducted extensive, unbiased AAMD simulations (18 independent trajectories, aggregation time ∼100 μs) starting from the TDP-43_310-350_ dimer models obtained from AF2-Multimer (**Fig. S2A,B**). These simulations, which provide atomic-level insights into helix-helix contacts, confirmed the dynamic behavior of TDP-43_310-350_ with dimers able to dissociate and re-associate within the microsecond timescale (**Fig. S2C**). The root-mean-square deviation of intermolecular atom distances (inter-dRMS) indicated highly fluctuating interhelical contacts (**Fig. S2D**).

The structural diversity within the dimer ensemble was characterized using the DSSP algorithm. The per-residue α-helical content averaged across 18 simulations showed >80% helicity for residues 320-331 and ∼30% for residues 339-343. Helicity in the CR region (aa: 320-331) increased by ∼20% in the dimeric state compared to the monomer (**Fig. S2E**), aligning with previous simulation results demonstrating helix stabilization upon dimerization^5^. Secondary structure maps highlighted diverse helical conformations with different helix lengths and positions (**Fig. S2F**). The most populated helical structures showed various intermolecular orientations, with both parallel and antiparallel binding modes (**Fig. S2F**). Using the simulated dimer ensemble, we characterized the intermolecular contacts stabilizing the helix-mediated dimerization. The dimer contacts (**Fig. 2D,E**) show significant differences compared to the initial contacts (**Fig. S2G**), highlighting dynamically interconverting structures.

We computed the average number of contacts between all possible residue type pairs in the TDP-43_310-350_ sequence (**Fig. S2H,I**). The total number of per-residue contacts revealed that W334, F313/316, and L330 form the most contacts (**Fig. 2F**), correlating with their role in stabilizing CTD condensates, whereas polar residues form fewer contacts, consistent with enhanced phase separation upon polar-to-alanine substitutions. We found a strong correlation (r=0.7) between the per-residue intermolecular contacts from dimer simulations and experimental phase separation (*c*_sat_) from alanine mutagenesis across CR (**Fig. 2G**). This support the idea that residues stabilizing CR helix-helix contacts in the dimer state contribute to the phase separation of CTD.

### TDP-43 CR helix-mediated assembly affects TDP-43 nuclear retention

Proper nuclear localization of TDP-43 is essential for its physiological function^34, 35^. The formation of higher-order assemblies through TDP-43 NTD and CTD self-interactions can be as important as RNA-binding via RRM for TDP-43 nuclear localization, as it prevents the passive diffusion of monomers to the cytoplasm by forming ribonucleoprotein complexes in the nucleus^36^. Disruption of RNA binding or self-assembly under abnormal conditions leads to increased levels of monomeric TDP-43, resulting in greater cytoplasmic localization. To understand how changes in CR helix-helix interactions and phase separation affect TDP-43 nuclear retention, we examined key alanine variants for their impact on cellular distribution^36^. These variants either greatly enhance (Q327A, Q331A, S333A, Q343A), moderately reduce (L340A), or impair (W334A and L330A) CTD condensation. We assessed the nuclear-cytoplasmic distribution of TDP-43 variants stably expressed in cell lines (HEK293)^3–6^. TDP-43 levels in nuclear and cytoplasmic fractions were quantified via immunoblotting (**Fig. S3**) and expressed as the nuclear-cytoplasmic (N/C) ratio (**Fig. 2H**).

W334A and L330A dramatically reduce the TDP-43 N/C ratio by approximately 90% and 80%, respectively, indicating strong depletion from the nucleus. This aligns with the effects of the deletion of CR^36^ or helix-disrupting mutant, A326P, which also disrupts TDP-43 nuclear localization (**Fig. 2H**). Conversely, variants that enhance helix-mediated self-assembly and phase separation (Q327A, Q331A, S333A, Q343A) also enhance nuclear localization, with S333A showing a 3-fold increase in nuclear retention compared to WT. Single glutamine-to-alanine substitutions generally increase the mean N/C ratio by about 50% compared to WT, although results for Q331A were not statistically significant. Unexpectedly, L340A increased nuclear TDP-43, contrary to our expectations based on its *c*_sat_. These findings suggest that CR-mediated assembly controls TDP-43 nuclear retention function, highlighting the need for further investigation into this mechanism.

### TDP-43 CTD forms higher-order oligomers

We used sedimentation velocity analytical ultracentrifugation (SV-AUC), an experimental tool that provides information on macromolecular size and shape in solution, to probe the size of TDP-43 CTD oligomers formed in solution. The distribution of sedimentation coefficients (corrected to standard conditions, water at 20°C), *s*_20,w_, at low concentrations of WT CTD show a peak at 1.5 S, corresponding to the monomeric CTD, and a second peak at 2.9 S at higher concentrations (**Fig. 3A**). These data suggest the presence of dimers and possibly larger species in equilibrium with monomers as concentration increases. To better characterize the assembled state, we experimentally trapped a dimeric state by forming a disulfide cross-link (S-S) between engineered cysteine residues at S273C (see Methods), as we previously used^5^. As expected, the apparent molecular weight of the cysteine cross-linked dimer is about twice that of the reduced monomeric control samples in nonreducing SDS-PAGE^5^ (see Methods). The cross-linked dimers also show the formation of a higher order assembled state with increasing concentration (**Fig. 3A**). However, the low solubility of S273C dimers still hinders the extensive characterization of the oligomer state.

**Figure 3.**
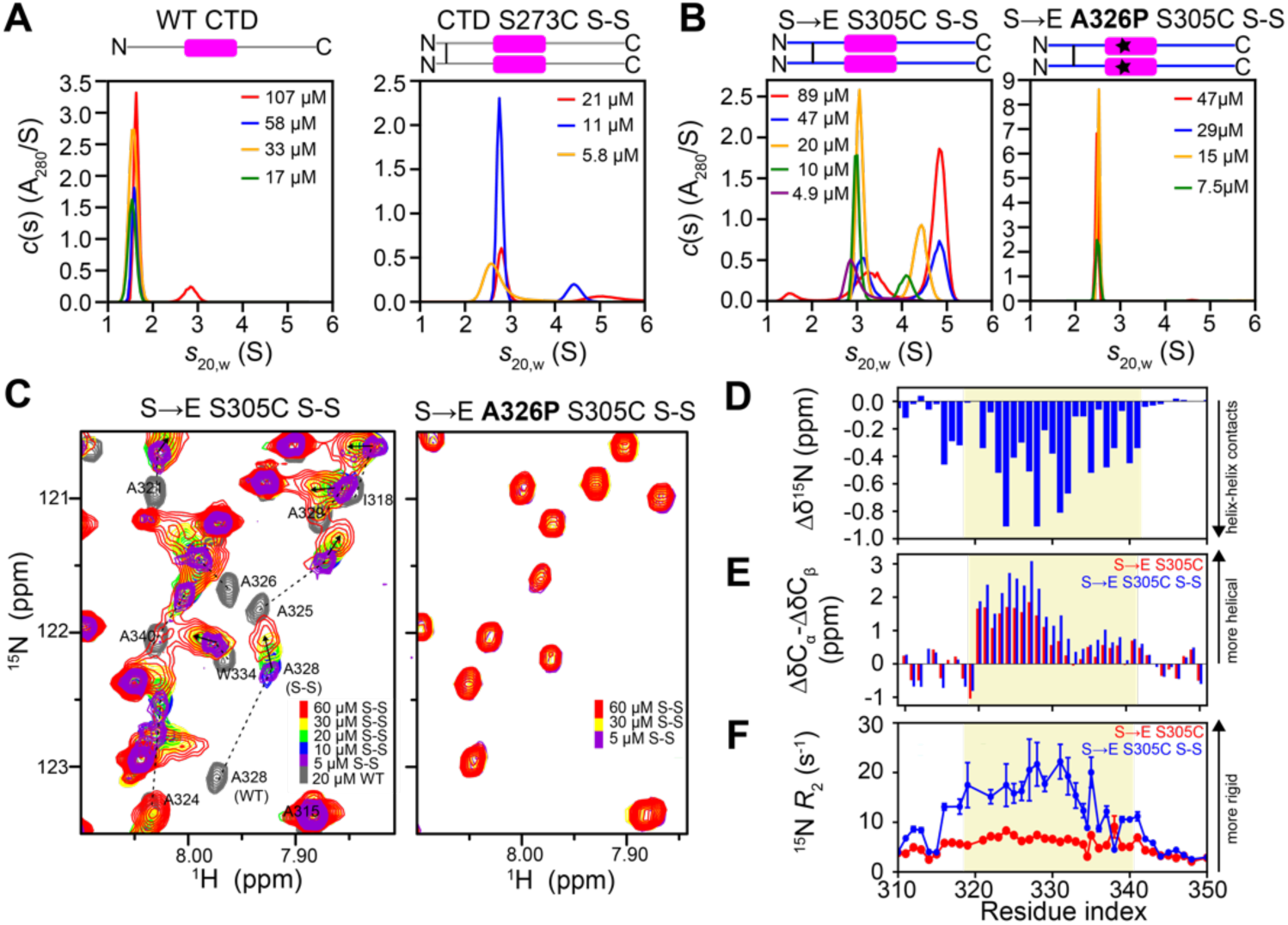
TDP-43 CTD forms higher-order oligomers larger than dimer, with increased local helicity and slowed motion. **A.** Sedimentation velocity analytical ultracentrifugation (SV-AUC) of WT CTD (without cross-linking) and CTD cysteine cross-linked dimers at S273C with increasing protein concentrations, at rotor speeds of 5000 rpm, 25°C. SV-AUC data were analyzed as a continuous *c*(s) distribution of sedimenting species. Sedimentation coefficients were corrected to standard conditions at 20 °C in water, *s*_20,w_. SV-AUC suggests the presence of species larger than dimer (∼2.4 S - 31.5 kDa). **B.** SV-AUC data as a function of protein concentration for S→E S305C variant show the presence of higher-order assembly (tetramer (∼3.6 S) and octamer (∼4.5 S)) larger than dimers (∼2.4 S), whereas only dimers are observed for the helix-disrupting S→E S305C A326P variant. **C.** NMR chemical shift analysis of S→E S305C as a function of increasing protein concentration exhibits distinct chemical shift changes distinct from dimer formation, suggesting formation of higher order oligomers with additional new contacts. Helix-disrupting mutant, A326P on S→E S305C dimer does not show chemical shift deviations up to 60 μM protein concentrations, suggesting the oligomerization is primarily mediated by helix-helix interaction. **D**. Quantification of ^15^N Δδ values between dimeric (cross-linked, oxidized) and monomeric (reduced with 1mM DTT) S→E S305C variants. The concentration of cross-linked S→E S305C is 30 μM, while monomeric S→E S305C (reduced) is 60 μM. Measurements performed in 20 mM MES buffer, pH 6.1 at 47°C. **E**. Experimental NMR secondary chemical shifts (ΔδC_α_-ΔδC_β_) of dimeric (cross-linked, oxidized) and monomeric (reduced) S→E S305C variants show the increase in helicity with cross-linking for entire CR residues. **F**. ^15^N *R*_2_ parameters for cross-linked (oxidized) and monomeric (reduced) S→E S305C variants suggest slowed reorientation motions for the 315–343 region upon cross-linking.

To overcome low solubility issues, we engineered a new CTD variant by introducing charged, glutamic acid residues at serine positions (S→E) in the disordered flanking regions of CTD, mimicking how phosphorylation may disrupt assembly^37, 38^ (See Table S1). This modification significantly enhanced solubility (> 100 μM) while preserving CR structure, as indicated by NMR spectral fingerprint (**Fig. S4**). To drive helical interactions, we designed two obligate dimers with cross-linking site closer to CR at position S305 and S317 (S→E S305C and S→E S317C). We modeled the cross-linked dimer, TDP-43_310–350 S317C S-S_, from AF2-Multimer predictions with a disulfide bond at S317C (**Fig. S5A**) and performed long, unbiased AAMD simulations (∼48 μs). The simulations showed that per-residue helix fractions and pairwise intermolecular contacts for the cross-linked dimer ensemble remained similar to WT dimer (**Fig. S5B,C**), with consistent major contributors to helix-helix contacts (W334, F316, L330) (**Fig. S5C**). This was further confirmed by a strong correlation between per-residue contacts in both cases (**Fig. S5D**), suggesting that the fundamental features of CR remain similar despite modifications to the flanking residues.

SV-AUC data of the solubilized S→E dimers also suggest the formation of high-order oligomers. Sedimentation profiles of S→E S273C dimers indicate mostly dimer species at all concentrations studied, with traces (<1%) of tetramer (**Fig. S6A**). Cross-linkage closer to the CR (S→E S305C S-S and S→E S317C) enhances the assembly. The sedimentation coefficient distributions for S→E S317C at different concentrations show the presence of two main peaks, corresponding to the dimer (∼2.4 S) and tetramer (∼3.6 S) states, with peaks of the latter are more pronounced at higher concentrations (**Fig. S6A**). The profiles for S→E S305C indicate peaks corresponding to dimer, tetramer, and octamer (∼4.5 S), with the fraction of the octamer increasing with protein concentration (**Fig. 3B**). Weighted-average sedimentation coefficient isotherms support a dimer-tetramer self-association for S→E S317C with a *K*_d_ of 46 µM, whereas the isotherm for S→E S305C supports a dimer–tetramer–octamer self-association with a dimer-tetramer *K*_d_ of 58 µM and a tetramer-octamer *K*_d_ of 4.7 µM (**Fig. S7A**). Both the sedimentation profiles (**Fig. S7B**) and the weighted average sedimentation coefficient isotherms indicate the dimer-tetramer self-association for S→E S317C, whereas cooperative self-assembly beyond tetramer is seen for S→E S305C. To demonstrate that higher-order assemblies are mediated by the helical region, we studied the oligomeric size distributions of the helix-breaking mutant, A326P on S→E S305C dimer. SV-AUC data shows the absence of the higher-order structures, with only peaks for a dimer state, suggesting that the helical region is essential for observed oligomerization (**Fig. 3B, Fig S7A**).

After confirming the presence of higher-order oligomers larger than dimer, we used NMR to characterize the structural changes upon helix-mediated TDP-43 CR oligomerization using the cross-linked S→E dimers. The 2D NMR spectrum of S→E dimers with cross-linking positions show chemical shift differences in the CR that increase as the cross-link is moved closer to the CR and lie along the same line formed by WT CTD suggesting transient formation of helix-helix dimers increases as the cross-linking distance to the CR is shortened (**Fig. 3C, S6B**). Large CSPs for CR residues as the protein concentration increases, especially for S305C and S317C, suggest multimerization beyond dimer occurs in this concentration range, consistent with AUC experiments. Importantly, these chemical shift perturbations lie along a new vector, consistent with the formation of a distinct multimeric structure (**Fig. 3C, S6B**)^16^. CR peaks broadened at higher concentrations, indicating the increasing population of the assembled state. Comparatively, the helix-breaking mutant, S→E A326P S305C dimer does not show CSPs up to 60 μM protein concentrations (**Fig. 3C**), confirming that higher-order oligomers, larger than dimer, observed via SV-AUC measurements, are mediated by TDP-43 CR.

### NMR structural characterization reveals the TDP-43 CR higher-order assembly with increased local helicity and rigidity

To probe structural changes upon CR-mediated oligomerization, we selected S→E S305C dimers for NMR structural measurements (at 47 °C to improve NMR spectra for CR residues) due to their tendency to form oligomers and better solubility. ^15^N chemical shift differences between the dimeric (oxidized) and monomeric (reduced) S305C variants showed upfield (negative) CSPs for residues 316-341 (**Fig. 3D**), consistent with enhanced helix-helix contacts and increased helicity^16, 39^. In the cross-linked S→E S305C dimers, pronounced positive increases in ^13^C secondary shifts were observed for the entire CR (**Fig. 3E**), suggesting enhanced helicity upon oligomerization in both the main helical region (aa: 320-331) and the subsequent region (aa: 332-343). ^15^N spin relaxation experiments revealed slowed backbone motions for the entire CR upon oligomerization with transverse relaxation rate constants (^15^N *R*_2_) showing around a four-fold increase compared to monomeric controls (**Fig. 3F**). This indicates slowed motions across residues 315–343 upon helix-helix assembly, consistent with the formation of locally rigid, higher-order structure within CR.

### NOEs show key contacts for TDP-43 higher order self-assembly

Next, we aimed to elucidate the structural details of the observed higher-order assembled states (larger than dimer), characterized by increased local helicity and rigidity using nuclear Overhauser effect (NOE) NMR. We conducted filtered, edited NOE-HSQC experiments 200 µM (1:1 mix of unlabeled and ^13^C/^15^N proteins) to selectively probe intermolecular contacts even in partially populated oligomeric assemblies^40^. For these measurements, we selected a highly soluble and stable TDP-43 CTD variant (without cross-links) that removes six phenylalanine residues in the IDRs flanking the CR (6F→A), preventing aggregation but preserving helix-helix interaction^20^. This construct is more suitable for NOE experiments compared to S→E S305C dimer, which tends to aggregate at the high concentrations required for the experiments. Experiments were performed at 12°C to enhance multimer population and NOEs by slowing molecular motion as well as to extend sample stability.

Two-dimensional NOE strips for NOEs to ^13^C-attached hydrogens suggest prevalent interactions involving aliphatic M/L/I/A methyl within the helical region (**Fig. 4A**), much more than control experiments using 100% labeled protein to prevent any intramolecular artifacts from incomplete labeling. NOEs are large for I318 but smaller for A315, consistent with structure starting forming primarily in the conserved region (**Fig. 4B**). NOEs to ^15^N attached hydrogens provide additional intermolecular interactions with residue-by-residue resolution, showing backbone and side chain amide hydrogens (NH and NH_2_) interact with methyl groups, aromatic and polar groups, and H_α_ (**Fig. 4C**). Importantly, NOEs to backbone NH hydrogens are observed for several positions from A325 to L340, suggesting structured interactions across the entire CR (**Fig. 4C**). Additonally, W334 sidechain positions interact with aliphatic methyl, aromatic, backbone amide, and H_α_ hydrogens (**Fig. 4C,D**). Contacts involving the phenyl group of phenylalanine (F313/316) with aliphatic methyl groups are also present (**Fig. 4D**), suggesting these residues contribute to multimerization.

**Figure 4.**
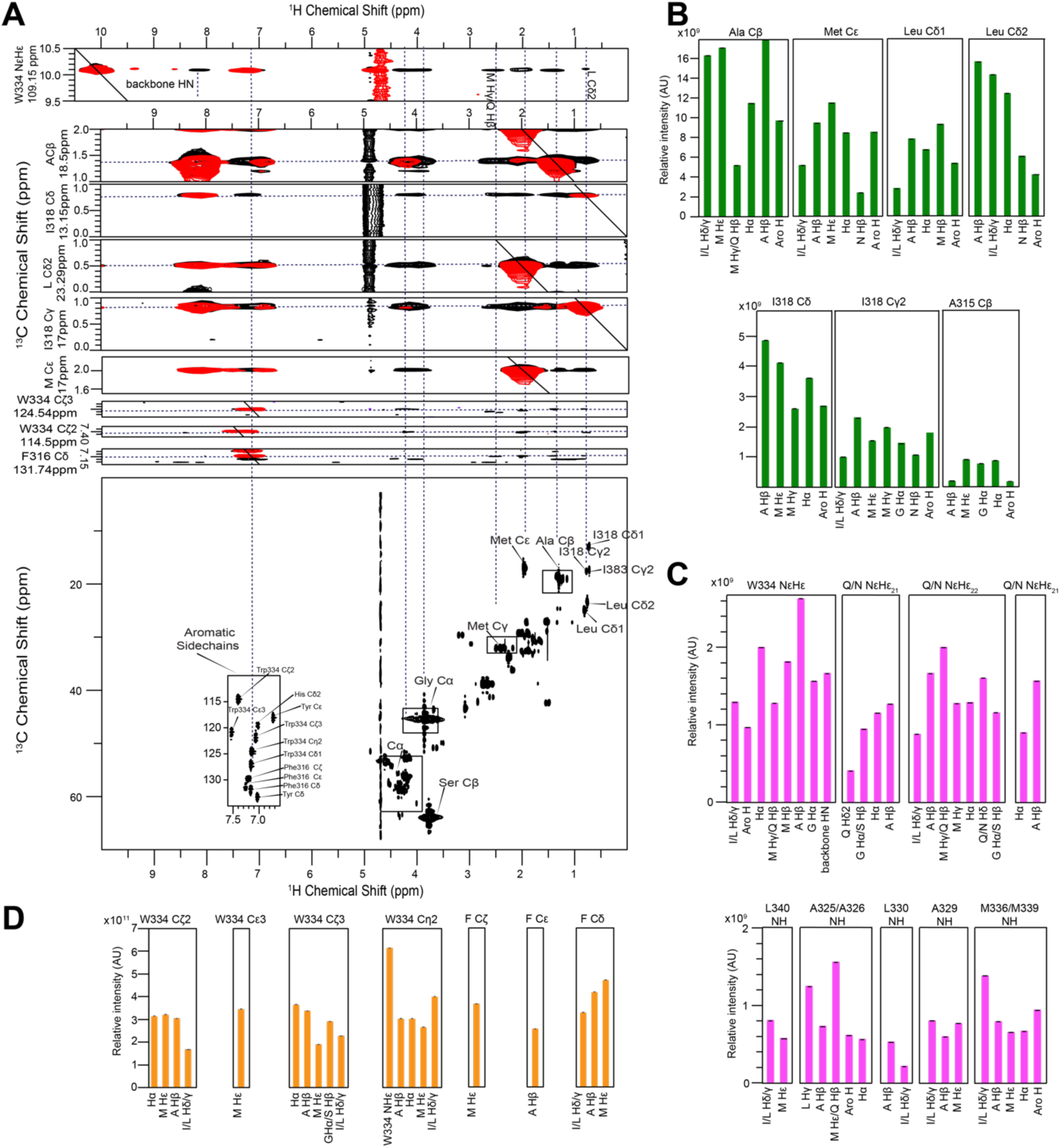
Intermolecular NOEs show key contacts for TDP-43 CR assembly. **A**. ^13^C filtered-edited NOE-HSQC strips of 200 μM 6F→A (1:1 mix of ^13^C/^15^N labeled and unlabeled, black) measured at 12 °C, 600 MHz, 20 mM MES pH 6.1, Transient NOEs (arising from equilibrium exchange between monomeric and multimeric states) are not artifacts as demonstrated by data from control sample (100% ^13^C/^15^N sample), red, that do not have extensive methyl NOEs. **B**. Quantification of NOEs from ^13^C-attached positions of different residues. Intensities were corrected for intramolecular artifacts arising from incomplete labeling by subtraction of 0.5*NOEs measured in a control sample 100% ^13^C/^15^N sample). **C**. Quantification of NOEs from ^15^N-attached positions of different residues. Intensities were corrected for artifacts as above. **D**. Quantification of NOEs from ^13^C-attached positions of aromatic residues. Intensities were corrected for artifacts as above.

Collectively, NOE experiments reveal the presence of contacts between many residue types in the TDP-43 CR assembly, prominently involving aliphatic and aromatic residues. However, the repetitive CR sequence (five M, six A, two S, three Q residues), relatively large separation between resolved backbone ^1^H/^15^N positions in adjacent helices, and difficulties in obtaining higher resolution/signal-to-noise data due to sample instability make it challenging to derive molecular models directly from these experimental constraints.

### AAMD simulations reveal the tetrameric helical assembly of TDP-43 CTD

To obtain an atomic structural model of TDP-43 CR, we turned to computer modeling using AF2-Multimer to predict potential oligomeric structures (trimer, tetramer, hexamer, octamer) of TDP-43_310-350_. Like for predicted dimers (see above), the model confidence score for all predicted multimeric structures is notably low (**Fig. 5A**), compared to a control homo-tetrameric coiled-coil sequence (CC-tet)^41^ (**Fig. 5A**). This suggests that AF2-Multimer cannot identify TDP-43 CR multimers or that the dynamic TDP-43 CR multimers^5^ are unlike stable coiled-coil sequences that are more easily predicted. Hence, we employed AAMD simulations (5 μs) to directly assess the structural stability of these oligomers and to explore whether AAMD could result in refined interhelical contacts that stabilize the assembled states (**Fig. 5B**).

**Figure 5.**
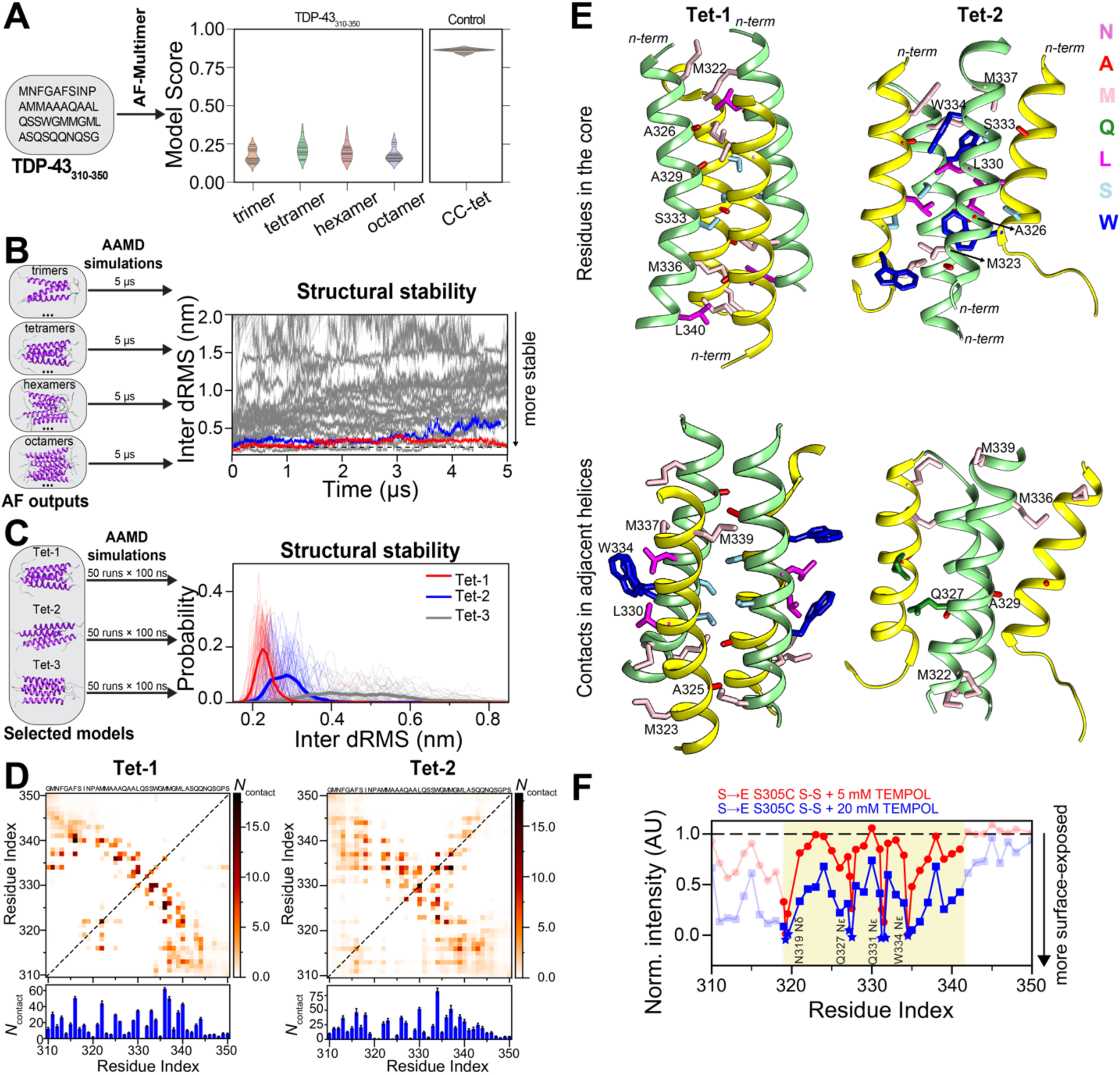
AAMD simulations highlight residues that stabilize tetrameric helical assembly of TDP-43 CTD. **A.** Potential oligomeric structures (trimer, tetramer, hexamer, octamer) of TDP-43_310-350_ fragment is predicted using AF2-Multimer. The model confidence score (0.8 × ipTM + 0.2 × pTM) of predicted structures for different multimeric states of TDP-43_310-350_ is compared against model confidence score of AF2-Multimer predicted tetrameric structures of homotetrameric coiled-coil sequence (CC-tet, 4-KE-4, PDB 6XY1), which served as a control. **B**. AF-2 Multimer predicted oligomeric structures is then fed to run unbiased AAMD simulations (5 μs each) and structural stability is analyzed by computing inter-dRMS of structures from AAMD simulation trajectories. Inter-dRMS of all heavy atoms (excluding hydrogen atoms) for residues from 320 to 341 as a function of time with respect to initial conformations are shown. (See Fig. S8 for more details). **C.** Inter-dRMS analysis from AAMD simulations (50 runs with 100 ns each) of selected tetramer models from the structural analysis shown in B. Inter-dRMS distribution of all heavy atoms (excluding hydrogen atoms) for residues from 320 to 341 as a function of time with respect to initial conformations are shown. The lighter colors represent the inter-dRMS distributions for 50 independent runs (100 ns each) for each model, while the darker colors represent the average inter-dRMS for Tet-1, Tet2 and Tet-3. Tet-1 exhibits greater stability compared to Tet-2 and Tet-3 models. **D.** (top) Pairwise intermolecular contact maps of Tet-1 and Tet-2 from AAMD simulations (single run, 5 μs). (bottom) Total number of contacts per residue position (N_contact_) derived through summation of all pairwise contacts along y-axis based on two dimensional pairwise intermolecular contact maps. **E.** Representative structures of the CR (aa:319-341) of Tet-1 and Tet-2 from the last microsecond of AAMD simulations (single run, 5 μs) are shown. Parallel helices are shown as yellow (chain 1 & 3) and light green (chain 2 & 4) colored ribbons. Side chains (excluding hydrogen atoms) of CR residues that are in the core (top) and in contact between adjacent helices (bottom) are shown as sticks, with their colors indicating different residue types as illustrated in the right corner. **F**. Solvent PRE experiments 30 μM S→E S305C cross-linked dimer with respect to different TEMPOL concentrations. Normalized intensity ratio at 5 and 20 mM TEMPOL concentrations are shown. Peak intensities were normalized based on the sample without TEMPOL.

Interestingly, all hexamer (five) and octamer (four) models are found to be less stable, showing the loss of intermolecular contacts in the binding interface compared to their initial conformation (**Fig. S8A**). Among the eight trimer models, one structure (Tri-1) with helices binding in a parallel orientation was found to be relatively stable (**Fig. S8A**), exhibiting well-preserved helicity and interfacial contacts in residues 320-333, while residues 334-341 lose their helicity (**Fig. S8B,C**). Among the tetramer structures, two tetramer structures (which we named Tet-1 and Tet-2) with initial four-fold symmetry where adjacent helices run antiparallel and have distinct register and contacts, exhibited greater stability during the simulation (**Fig. S8A**). To further validate Tet-1 and Tet-2 stability, we performed 50 independent AAMD simulations (each 100 ns) of Tet-1 and Tet-2, along with Tet-3 as a control, which was found to be unstable in longer simulations (**Fig. 5B**). The inter-dRMS distributions from these simulations showed higher stability for Tet-1, followed by Tet-2, compared to Tet-3 (**Fig. 5C**). Collectively, AAMD simulations of AF-predicted oligomers reveal at least two stable tetramer structures, suggesting that dynamic nature of CR self-interactions may result in long-lived metastable configurations. A two-dimensional intermolecular pairwise contact map reveals that in both tetramer structures, adjacent helices primarily exhibit antiparallel contacts, with parallel helix-helix interactions occurring to a lesser extent between non-adjacent helix pairs (**Fig. 5D**). The core of the Tet-1 is composed of residues M322, A326, A329, S333, M336 and L340, which stabilize the assembly by forming the highest total number of contacts (**Fig. 5D,E**). Importantly, these residues correspond closely to the positions that give rise to intermolecular backbone HN NOEs, suggesting they are at the core in the experimental multimers. The stability is further enhanced by interhelical contacts between adjacent helices involving residues M323, A325, L330, S332, W334, M337 and M339 (**Fig. 5D,E**). Notably, W334 and L330 form an extended hydrophobic patch, enhancing the overall stability of the complex (**Fig. 5E**). In Tet-2, which settles into a stable state in AAMD simulation, the initial four-fold symmetry from the initial AF-based structure is lost due to partly destabilization of one helical unit (**Fig. 5E**). The interfacial contacts of the resulting tetrameric structure are different from those in Tet-1, being formed by M323, A326, L330, S333, W334, and M337, while M322, Q327, A329, M336, and M339 form interhelical contacts between adjacent helices. W334-L330 contacts are also observed in Tet-2, suggesting their critical role in tetramer formation. Interestingly, W334 which was in the incompletely packed core of the Tet-2 initial structure, starts to be surface-exposed as in Tet-1. Importantly, the intermolecular contacts in these tetramers are qualitatively consistent with experimental NOEs, showing interactions primarily from methyl groups and backbone NH atoms of aliphatic residues (M/A/L/I), W334 with methyl groups (M/A/L/I), and polar residues (Q/N/S).

We next assessed whether these structures agree with our NMR-derived measurements. The simulated transverse relaxation rate, ^15^N *R*_2_, of the Tet-1 and Tet-2 models demonstrate higher values compared to monomeric and dimeric ensembles (**Fig. S8D**), consistent with the slowed motions observed in experiments where the multimers are partially populated (**Fig. 3F**). The helix fraction computed from atomistic simulation of Tet-1 and Tet-2 using DSSP^42^ also aligns qualitatively with experimental results (**Fig. S8E**).

Importantly, both models exhibit numerous contacts and stable secondary structure of the region spanning G335-Q343, suggesting a reason for the perfect conservation in vertebrates of this region despite being unstructured in the monomeric form^16^. Structure in this region is also absent in the trimer models, further supporting a tetrameric structure of TDP-43 CR (**Fig. S8E**). Furthermore, structure in this region in the tetramer models explains the disruption of phase separation caused by ALS-associated M337V mutant and engineered variants M337P^16^ and M337A.

To further assess the relevance of the structural models, we performed solvent paramagnetic relaxation enhancement (sPRE) measurements (**Fig. S9**). This NMR-based method qualitatively characterizes the relative solvent accessibility of residues within biomolecular structures^43^. The sPRE pattern for cross-linked dimeric variants of S→E S305C showed some residues are more protected. Cross-linked dimers revealed complete signal loss for the sidechain NH groups of W334, Q331, Q327, and N319 (**Fig. 5F**), suggesting these groups are highly solvent exposed even upon multimerization. Indeed, our structural models show that the side chains of N319, Q327, Q331, and W334 are more solvent-exposed than other residues (**Fig. 5E**). Together, the models for the TDP-43 CR tetramer, exhibiting increased local helicity and rigidity, qualitatively agree with the determined surface accessible positions from sPRE experiments and NOEs on intermolecular interactions.

## Conclusions

Uncovering the atomic structural details of TDP-43 CR in its self-assembly pathway is crucial for a mechanistic understanding of CTD functional states and their disruption in disease. In this study, we definitively establish α-helical oligomers as a native form of the CR which is important for its splicing and nuclear retention functions.^5, 6, 36^ Although our previous work showed that the first half of the CR (aa:320-330) was partially structured in the monomeric form^16^, these results explain why the entire 21-residue CR is conserved – to mediate assembly. Unlike many known coiled-coils and leucine zippers, the functional CR-mediated assembly is characterized by dynamic and low affinity α- helical interactions, likely adopting multiple stable configurations with methionine side chains at the core of the assembly. These dynamic CR self-interactions are commensurate in affinity with the transient, multivalent contacts formed by the disordered regions^20^, N-terminal domain^13^, and RRMs^14, 44^ of TDP-43^45^. Importantly, our integrative approach in determining atomic structural models of TDP-43 oligomeric state also shows that although AI-based predictors cannot currently discriminate between the possible conformers of dynamic assemblies, pairing these predictors with molecular simulation and experiment allows for refinement of the structural models that may serve to improve future predictions as more data on dynamic assemblies become available.

The sensitivity of TDP-43 nuclear retention to single CR mutations that alter CTD self-assembly highlight the precise tuning of TDP-43 assembly and function. Hence, a series of TDP-43 CR variants may serve as a way to precisely control its nuclear concentration and function^5^. Additionally, our structure model rationalizes the impact of disease mutations, such as M337V, that disrupt the CR assembly without altering the monomeric structure^16^. Even without explicit NLS-disrupting variants, as seen in cases of familial ALS of the related protein FUS, TDP-43 CTD mutants that disrupt helical assembly may drive TDP-43 aggregation by increasing TDP-43 cytoplasmic accumulation. These data also suggest that caution should be exercised in interpretation of localization and function of TDP-43 bearing protein domain tags which may influence its size and hence the nuclear/cytoplasmic equilibrium.

Our data support a model in which significant conformational rearrangement must occur for the formation of the β-sheet assemblies of TDP-43 CTD, including the CR, in ALS and neurodegeneration^9, 10^. Our recent work suggests that the CR self-interactions are essential not only for TDP-43 function but also for its aggregation^23^. Given that the CR must convert to β-sheet aggregates to form pathogenic fibrils^9, 10, 46^, an intriguing therapeutic avenue may be to pharmacologically stabilize the helical assembly observed here to prevent conversion of TDP-43 CTD into β-sheet aggregates. However, as TDP-43 participates in neuronal transport granules^47^, it is unclear if excessive nuclear retention of TDP-43 will result in toxic loss of its cytoplasmic function, another important avenue to explore in future cellular studies. Prior work has also suggested the Q/N-rich region directly following the CR may contribute to CTD aggregation^48^. Our simulations suggest this region can also nucleate transient intermolecular β-sheets between residues flanking the CR in the tetrameric helical assemblies (**Fig. S10**), which could then spread and lead to conversion of the CR helix into β-sheet structures. Hence, the assembly of TDP-43 CR appears delicately balanced to drive precisely tuned functional assembly^5^ but discourage aggregation, making it a challenging future drug target.

## Materials and Methods

### Expression and purification of recombinant proteins

All TDP-43 CTD variants were produced using codon-optimized sequences from a pJ411 bacterial expression vector as previously^16^. The expression was carried out in BL21 Star (DE3) *Escherichia coli* cells obtained from Life Technologies. The proteins were expressed in either LB or M9 minimal media supplemented with ^15^NH_4_Cl, following a slightly modified version of previously described as follows^5, 16^. Bacterial cultures were induced at an optical density (OD) of 0.8 with 1 mM IPTG and incubated for 4 hours at 37°C and 220 rpm. The cells were then collected by centrifugation (6000 rpm, 15 minutes, 4°C). The resulting cell pellets (2 liters of culture) were resuspended in a 20 mL buffer (20 mM Tris, 500 mM NaCl, 10 mM imidazole, pH 8.0), lysed using an ultrasonic cell disrupter, and the lysate was cleared by centrifugation (15,000 g, 1 hour, 4°C). The insoluble material, containing the inclusion bodies, was resuspended in a 40 mL solubilizing buffer (8 M urea, 20 mM Tris, 500 mM NaCl, 10 mM imidazole, pH 8.0) and the cell debris was cleared by centrifugation (25,000 rpm, 1 hour, 19°C).

The supernatant was then filtered using a 0.45 μm syringe filter, and the protein was purified using a 5 mL Histrap HP column with a gradient of 10 to 500 mM imidazole added to the solubilizing buffer. The purified protein fractions were desalted using a HiPrep 26/60 Desalting Column into TEV cleavage buffer (20 mM Tris, 500 mM GdnHCl, pH 8.0) and subjected to TEV cleavage overnight at room temperature. After cleavage, solid urea was added to the solution to reach a concentration of approximately 8 M urea. The resolubilized protein was then applied to the Histrap HP column to remove the histidine tag and histidine-tagged TEV protease. The cleaved protein fractions were concentrated, buffer-exchanged into a storage buffer (20 mM MES, 8 M urea, pH 6.1) at approximately 1.5 mM, aliquoted, flash-frozen, and stored at −80°C for further use. Samples with disulfide cross-links were created using copper phenanthroline catalysis and purified to ensure removal of residual (non-crosslinked) monomers by size exclusion chromatography, as previously described^5^.

### Microscopy

Following the manufacturer’s instructions, the protein stocks in 8 M urea were diluted 8x (to 1 M urea) with experimental buffer (20 mM MES, pH 6.1) to a final concentration of 150 to 200 μM TDP-43 CTD and then buffer exchanged into experimental buffer with equilibrated 0.5 mL Zeba spin desalting columns from Thermo Scientific following manufacturer instructions. An equal volume of NaCl stock solution prepared in MES buffer was added to obtain final salt concentration of 150 mM to induce phase separation. The samples were gently mixed, and the phase separation was monitored using DIC micrographs obtained with a Nikon Ti2-E Fluorescence Microscope equipped with a 40x objective. To capture the images, 10 μL of each sample was spotted onto a coverslip. The resulting images were subsequently processed using Fiji software.

### In vitro phase separation assay for determining saturation concentration

To quantitatively analyze the phase separation of CTD mutants, we performed assays to determine the saturation concentration by measuring the protein concentration in the supernatant after centrifuging samples with increasing salt concentrations, which induced phase separation. During centrifugation, the protein in micrometer-sized droplets sediments, representing the phase-separated state. The protein remaining in the supernatant corresponds to the dispersed phase. By measuring the amount of protein remaining in the supernatant, we determined the saturation concentration, *c*_sat_, which is the concentration above which the protein undergoes phase separation at the given condition. This value decreases with higher salt concentrations. To initiate phase separation, desalted protein samples were diluted to 80 μM using MES buffer. An equal volume of NaCl stock solution prepared in MES buffer was added to achieve final salt concentrations of 0, 37.5, 75, 150, and 300 NaCl. The samples were gently mixed and then centrifuged for 10 minutes at 12,000 rpm at room temperature. After centrifugation, the protein concentration in the supernatant was measured using a Nanodrop 2000c spectrophotometer. All measurements were performed in triplicate to ensure the accuracy and consistency of the data points.

### NMR spectroscopy

NMR spectroscopy was conducted using Bruker Avance 850 MHz or 600 MHz ^1^H Larmor frequency spectrometers equipped with HCN TCI z-gradient cryoprobes at a temperature of 320K or 298K. The NMR samples were prepared in 20 mM MES buffer at pH 6.1, with the addition of 5% ^2^H_2_O to serve as a lock solvent. The NMR sample in Figure 1 for the native condition was prepared in 20 mM HEPES, 150 mM NaCl buffer pH 7 with the addition of 5% ^2^H_2_O to serve as a lock solvent. Backbone assignments for the dimers were obtained through standard triple resonance experiments (HNCACB, HNCA, and CBCACONH) using Bruker pulse sequences. The acquired NMR data were processed using Bruker Topspin software and analyzed using CCPN^49^. Detailed information on the experimental NMR pulse sequences and parameters can be found in our previous publications^5, 16^. The ^15^N spin relaxation experiments at 850 MHz were recorded with 128 and 4096 total points, with acquisition times of 39 ms and 200 ms, in the indirect ^15^N and direct ^1^H dimensions, respectively. The sweep width was set to 19 ppm in the indirect ^15^N dimension and 12 ppm in the direct ^1^H dimension, centered at 116.6 ppm and 4.7 ppm. The ^15^N *R*_2_ experiments consisted of six interleaved relaxation delays, with an interscan delay of 2.5 s. The CPMG field was set to 556 Hz, and the total *R*_2_ relaxation CPMG loop lengths were 16.5 ms, 264.4 ms, 33.1 ms, 132.2 ms, 66.1 ms, and 198.3 ms. Each ^15^N *R*_1_ experiment consisted of six interleaved ^15^N *R*_1_ relaxation delays: 5 ms, 1,000 ms, 100 ms, 800 ms, 500 ms, and 300 ms. The (^1^H) ^15^N heteronuclear NOE experiments were conducted using interleaved sequences with and without proton saturation and a recycle delay of 5 s. The (^1^H) ^15^N hetNOE experiments were recorded with 256 and 4,096 total points in the indirect ^15^N and direct ^1^H dimensions, respectively. Additional information on the experimental relaxation NMR parameters can be found in our previous publication^50^.

For the PRE experiments, the TEMPOL reagent was purchased from Sigma-Aldrich. The protein stocks were diluted from storage buffer containing 8 M urea and buffer exchange to 20 mM MES at pH 6.1, followed by the addition of 1.0 M TEMPOL stock solutions to achieve final concentrations of 0, 5, and 20 mM TEMPOL in a 30 μM (in dimer units) protein solution. Subsequently, ^1^H-^15^N HSQC experiments were conducted on an 850 MHz spectrometer with acquisition parameters set at 4096 direct points and 256 indirect points. The acquired NMR data were processed using NMRPIPE with consistent parameters, and peak intensities were analyzed using CCPN software. Peak intensities were normalized to intensities of samples without TEMPOL to determine the attenuation of resonances induced by the presence of TEMPOL.

### Analytical ultracentrifugation

Sedimentation velocity experiments were conducted at the specified temperature on a Beckman Coulter ProteomeLab XL-I analytical ultracentrifuge. 400 µL of each of the samples were loaded into prechilled 2-channel centerpiece cells and rotor, allowed to equilibrate at the temperature specified under vacuum for approximately 20 hours and then analyzed at either 50 krpm over a period of 24 - 27 hours. Data were collected using interference optical systems with scans collected at 7 minute intervals. Data were analyzed in SEDFIT 12.4434 in terms a continuous *c*(s) distribution of Lamm equation solutions using an uncorrected s range of 0.0 – 10.0 S with a resolution of 200 and a confidence level of 0.68. When necessary, early timepoint scans were not used for model fitting to remove contributions from large aggregates. Solution densities ρ were measured at the specified temperature on an Anton Paar DMA 5000 density meter and solution viscosities η were measured at the specified temperature using an Anton Paar AMVn rolling ball viscometer. The partial specific volumes v of the peptides were calculated in SEDNTERP 1.0935, and corrected for isotopic substitution. Sedimentation coefficients s were corrected to s_20,w_.

### Cloning, cell culture, and stable cell production

Generation of stable cells lines HEK293^HA-TDP-43^ expressing HA-tagged WT, and A326P was previously described^36^. Constructs to express TDP-43 mutants Q327A, L330A, Q331A, S333A, W334A, L340A, and Q343A were generated by site-directed mutagenesis as previously described^3–6^ using the primers listed in TABLE S2. Generation of HEK293^HA-TDP-43^ stable cell lines, expressing WT and mutant HA-tagged TDP-43 was carried out according to previously described protocols^14, 15^.

### TDP-43 in-cell nuclear localization experiments

In brief, HEK293-Flp-In T-Rex 293 cells (Thermo Fisher Scientific) were stably transfected to express HA-TDP-43 upon induction with tetracycline (1 μg/ml). Cells were grown and maintained in DMEM (Dulbecco’s Modified Eagle’s Medium–High Glucose, Corning) supplemented with 10% FBS (fetal bovine serum) and incubated in a humid atmosphere at 37°C and 5% CO_2_. Expression of HA-tagged TDP-43 construct was induced at 30% confluence for 48 h. Cell fractionation, immunoblotting, and quantification of nuclear to cytoplasmic ratio of HA-TDP-43 variants was performed similarly to previously described protocols^36^, probing with TDP-43 antibody (10782-2-AP, Proteintech).

### AlphaFold-Multimer Predictions

TDP-43_310-350_ oligomers were generated using AlphaFold2-Multimer, providing a multi-sequence FASTA file of TDP-43_310-350_. The predictions for multimers were executed based on multiple sequence alignment (MSA) alone, with the exclusion of PDB templates through the use of the parameter --max_template_date=1950-1-1. The full database (-- db_preset=full_dbs) were used to create the alignments. Dimer and tetramer models were generated utilizing both AlphaFold v2.2.0 and v2.3.0, while trimer, hexamer, and octamer structures were generated using AlphaFold v2.3.0. In AlphaFold-Multimer v2.2.0, one seed per model was employed, resulting in a total of five predictions. In AlphaFold-Multimer v2.3.0, five seeds were used per model, yielding a total of 25 predictions. The relaxed structures (--use_gpu_relax=True) without major clashes were employed as the initial configurations in AAMD simulations. It is important to note that the predicted structures contain major clashes, especially in the case of higher-order oligomers; thus, the number of structures simulated for each oligomeric state differs from one another.

### All-atom MD simulations

#### Initial structure, force-field choise and system setup

The monomer structures (five models) of TDP-43_310-350_ from AlphaFold2 (AF2) and the dimer (5 models from v2.2.0 and 13 models from v2.3.0), trimer (8 model from v2.3.0), tetramer (2 models from v2.2.0 and 5 models from v2.3.0), hexamer (5 models from v2.3.0), and octamer (4 models from v2.3.0) structures of TDP-43_310-350_ from AlphaFold2-Multimer were employed as the starting configurations for AAMD simulations with explicit solvent and ions. The protein was modelled using the Amber03ws force field^51^ (https://bitbucket.org/jeetain/all-atom_ff_refinements/src/master/) and solvated with TIP4P/2005 water model^52^. To mimic the physiological salt concentration (150 mM), Na^+^ and Cl^−^ ions were added to the protein-water system. The improved salt parameters from Lou and Roux were used for all simulations^53^. The protein was solvated in a truncated octahedron box, ensuring 1.2 nm separation between protein atoms and box edges.

#### Simulation protocol

Following solvation, the system was minimized via the steepest descent algorithm using GROMACS 2020^54^. After minimization, the Nose-Hoover thermostat^55^ was used for temperature equilibration at 300 K with a coupling constant of 1 ps for protein, water and ions. After the NVT equilibration, a 100 ns NPT equilibration was run using the Berendsen barostat^56^ with isotropic coupling and a coupling constant of 5 ps to achieve pressure of 1 bar. Production simulations were conducted in the NPT ensemble (1 bar, 300 K) using the Langevin Middle Integrator^57^ (friction coefficient = 1 ps^-1^) and the Monte Carlo Barostat with isotropic coupling in AMBER 22^58^. Hydrogen mass was increased to 1.5 amu to enable a timestep of 4 fs. Short-range nonbonded interactions were calculated based on a cutoff radius of 0.9 nm, while long-range electrostatics were handled with the Particle Mesh Ewald (PME) method^59^. Hydrogen-related bonds were constrained using the SHAKE algorithm^60^.

#### Trajectory analysis

In contact analysis, two residues *i* and *j* with sequence separation greater than 3 (|*i*-*j*|>3) were counted in contact if any two heavy atoms of the residues were within 0.6 nm of each other. This cutoff has been extensively used in our previous studies and has been thoroughly explained in the work of Zheng et al.^61^ Helix fraction calculations were performed using the gmx do_dssp, relying on the DSSP library^42^. Minimum distance between the monomeric units in the two-chain simulation trajectories were computed using gmx mindist. The distance root mean square deviation of intermolecular atom distances (inter-dRMS) for all heavy atoms across the CR (amino acids 320-341) from reference structures was calculated in PLUMED^62^ using the DRMSD command. The NMR relaxation parameters (^15^N *R*_2_) were computed using a previously published method^16, 63^. The most representative structures from AAMD simulations of oligomer models were determined using the gmx cluster tool with a GROMOS clustering algorithm. Backbone atoms of CR residues (aa: 320-341) were considered, with an RMSD cutoff of 0.3 nm, based on the last microsecond of the trajectory. All snapshots from atomistic simulations were generated using UCSF Chimera^64^/ChimeraX^65^.

## Acknowledgements

This research was supported in part by NINDS and NIA R01NS116176 (to NLF and JM). JS was supported by a Milton Safenowitz Postdoctoral Fellowship from the ALS Association and Judith and Jean Pape Adams Postdoctoral Fellowship at Brown University. RG was supported by the Intramural Research Program of the NIH, The National Institute of Diabetes and Digestive and Kidney Diseases (NIDDK). NMR experiments were conducted with the support of the Structural Biology Core Facility in the Division of Biology and Medicine at Brown University and with assistance from Mandar Naik. Use of the Texas A&M High Performance Research Computing is greatly acknowledged for the computational resources utilized in this work. The funders had no role in study design, data collection and analysis, decision to publish or preparation of the manuscript.

## References

1 Chen-Plotkin, A. S., Lee, V. M. Y. & Trojanowski, J. Q. TAR DNA-binding protein 43 in neurodegenerative disease. Nature Reviews Neurology 6, 211–220 (2010). 10.1038/nrneurol.2010.18

2 Cohen, T. J., Lee, V. M. & Trojanowski, J. Q. TDP-43 functions and pathogenic mechanisms implicated in TDP-43 proteinopathies. Trends in molecular medicine 17, 659–667 (2011).

3 Hallegger, M. et al. TDP-43 condensation properties specify its RNA-binding and regulatory repertoire. Cell 184, 4680–4696. e4622 (2021).

4 Tziortzouda, P., Van Den Bosch, L. & Hirth, F. Triad of TDP43 control in neurodegeneration: autoregulation, localization and aggregation. Nature Reviews Neuroscience 22, 197–208 (2021). 10.1038/s41583-021-00431-1

5 Conicella, A. E. et al. TDP-43 α-helical structure tunes liquid–liquid phase separation and function. Proceedings of the National Academy of Sciences 117, 5883–5894 (2020). 10.1073/pnas.1912055117

6 Koehler, L. C. et al. TDP-43 oligomerization and phase separation properties are necessary for autoregulation. Frontiers in Neuroscience 16, 818655 (2022).

7 Neumann, M. et al. Ubiquitinated TDP-43 in frontotemporal lobar degeneration and amyotrophic lateral sclerosis. Science 314, 130–133 (2006).

8 Li, H.-Y., Yeh, P.-A., Chiu, H.-C., Tang, C.-Y. & Tu, B. P.-h. Hyperphosphorylation as a defense mechanism to reduce TDP-43 aggregation. PloS one 6, e23075 (2011).

9 Arseni, D. et al. TDP-43 forms amyloid filaments with a distinct fold in type A FTLD-TDP. Nature 620, 898–903 (2023). 10.1038/s41586-023-06405-w

10 Arseni, D. et al. Structure of pathological TDP-43 filaments from ALS with FTLD. Nature 601, 139–143 (2022). 10.1038/s41586-021-04199-3

11 Nelson, P. T. et al. Limbic-predominant age-related TDP-43 encephalopathy (LATE): consensus working group report. Brain 142, 1503–1527 (2019).

12 Afroz, T. et al. Functional and dynamic polymerization of the ALS-linked protein TDP-43 antagonizes its pathologic aggregation. Nature Communications 8, 45 (2017). 10.1038/s41467-017-00062-0

13 Wang, A. et al. A single N-terminal phosphomimic disrupts TDP-43 polymerization, phase separation, and RNA splicing. EMBO J 37 (2018). 10.15252/embj.201797452

14 Lukavsky, P. J. et al. Molecular basis of UG-rich RNA recognition by the human splicing factor TDP-43. Nature Structural & Molecular Biology 20, 1443–1449 (2013). 10.1038/nsmb.2698

15 Grese, Z. R. et al. Specific RNA interactions promote TDP-43 multivalent phase separation and maintain liquid properties. EMBO reports 22, e53632 (2021).

16 Conicella, Alexander E., Zerze, Gül H., Mittal, J. & Fawzi, Nicolas L. ALS Mutations Disrupt Phase Separation Mediated by α-Helical Structure in the TDP-43 Low-Complexity C-Terminal Domain. Structure 24, 1537–1549 (2016). 10.1016/j.str.2016.07.007

17 King, O. D., Gitler, A. D. & Shorter, J. The tip of the iceberg: RNA-binding proteins with prion-like domains in neurodegenerative disease. Brain research 1462, 61–80 (2012).

18 Johnson, B. S. et al. TDP-43 is intrinsically aggregation-prone, and amyotrophic lateral sclerosis-linked mutations accelerate aggregation and increase toxicity. Journal of Biological Chemistry 284, 20329–20339 (2009).

19 Buratti, E. Functional significance of TDP-43 mutations in disease. Advances in genetics 91, 1–53 (2015).

20 Mohanty, P. et al. A synergy between site-specific and transient interactions drives the phase separation of a disordered, low-complexity domain. Proceedings of the National Academy of Sciences 120, e2305625120 (2023).

21 Schmidt, H. B., Barreau, A. & Rohatgi, R. Phase separation-deficient TDP43 remains functional in splicing. Nature Communications 10, 4890 (2019). 10.1038/s41467-019-12740-2

22 Lim, L. Z., Wei, Y. Y., Lu, Y. M. & Song, J. X. ALS-Causing Mutations Significantly Perturb the Self-Assembly and Interaction with Nucleic Acid of the Intrinsically Disordered Prion-Like Domain of TDP-43. Plos Biol 14 (2016). e1002338. 10.1371/journal.pbio.1002338

23 Xiao, Y. et al. Intra-condensate demixing of TDP-43 inside stress granules generates pathological aggregates. bioRxiv, 2024.2001.2023.576837 (2024). 10.1101/2024.01.23.576837

24 Sharma, K. et al. Cryo-EM observation of the amyloid key structure of polymorphic TDP-43 amyloid fibrils. Nature Communications 15, 486 (2024). 10.1038/s41467-023-44489-0

25 Jumper, J. et al. Highly accurate protein structure prediction with AlphaFold. Nature 596, 583–589 (2021). 10.1038/s41586-021-03819-2

26 Richard, E. et al. Protein complex prediction with AlphaFold-Multimer. bioRxiv, 2021.2010.2004.463034 (2022). 10.1101/2021.10.04.463034

27 Bai, Y., Milne, J. S., Mayne, L. & Englander, S. W. Primary structure effects on peptide group hydrogen exchange. *Proteins: Structure*, Function, and Bioinformatics 17, 75–86 (1993). 10.1002/prot.340170110

28 Kjaergaard, M., Brander, S. & Poulsen, F. M. Random coil chemical shift for intrinsically disordered proteins: effects of temperature and pH. Journal of Biomolecular NMR 49, 139–149 (2011). 10.1007/s10858-011-9472-x

29 Kjaergaard, M. & Poulsen, F. M. Sequence correction of random coil chemical shifts: correlation between neighbor correction factors and changes in the Ramachandran distribution. Journal of Biomolecular NMR 50, 157–165 (2011). 10.1007/s10858-011-9508-2

30 Zerze, G. H., Zheng, W., Best, R. B. & Mittal, J. Evolution of All-Atom Protein Force Fields to Improve Local and Global Properties. The Journal of Physical Chemistry Letters 10, 2227–2234 (2019). 10.1021/acs.jpclett.9b00850

31 Fonda, B. D., Jami, K. M., Boulos, N. R. & Murray, D. T. Identification of the Rigid Core for Aged Liquid Droplets of an RNA-Binding Protein Low Complexity Domain. Journal of the American Chemical Society 143, 6657–6668 (2021). 10.1021/jacs.1c02424

32 Shenoy, J. et al. Structural dissection of amyloid aggregates of TDP-43 and its C-terminal fragments TDP-35 and TDP-16. The FEBS Journal 287, 2449–2467 (2020). 10.1111/febs.15159

33 Shenoy, J. et al. Structural polymorphism of the low-complexity C-terminal domain of TDP-43 amyloid aggregates revealed by solid-state NMR. Frontiers in Molecular Biosciences 10, 1148302 (2023).

34 Tollervey, J. R. et al. Characterizing the RNA targets and position-dependent splicing regulation by TDP-43. Nature neuroscience 14, 452–458 (2011).

35 Polymenidou, M. et al. Long pre-mRNA depletion and RNA missplicing contribute to neuronal vulnerability from loss of TDP-43. Nature neuroscience 14, 459–468 (2011).

36 dos Passos, P. M., Hemamali, E. H., Mamede, L. D., Hayes, L. R. & Ayala, Y. M. RNA-mediated ribonucleoprotein assembly controls TDP-43 nuclear retention. Plos Biol 22, e3002527 (2024). 10.1371/journal.pbio.3002527

37 Gruijs da Silva, L. A., et al. Disease-linked TDP-43 hyperphosphorylation suppresses TDP-43 condensation and aggregation. EMBO J 41, e108443 (2022). 10.15252/embj.2021108443

38 Monahan, Z. et al. Phosphorylation of the FUS low-complexity domain disrupts phase separation, aggregation, and toxicity. EMBO J 36, 2951–2967 (2017). 10.15252/embj.201696394

39 Wang, Y. & Jardetzky, O. Probability-based protein secondary structure identification using combined NMR chemical-shift data. Protein Science 11, 852–861 (2002). 10.1110/ps.3180102

40 Zwahlen, C. et al. Methods for measurement of intermolecular NOEs by multinuclear NMR spectroscopy: application to a bacteriophage λ N-peptide/boxB RNA complex. Journal of the American Chemical Society 119, 6711–6721 (1997).

41 Edgell, C. L., Savery, N. J. & Woolfson, D. N. Robust De Novo-Designed Homotetrameric Coiled Coils. Biochemistry 59, 1087–1092 (2020). 10.1021/acs.biochem.0c00082

42 Kabsch, W. & Sander, C. Dictionary of protein secondary structure: Pattern recognition of hydrogen-bonded and geometrical features. Biopolymers 22, 2577–2637 (1983). 10.1002/bip.360221211

43 Clore, G. M. & Iwahara, J. Theory, Practice, and Applications of Paramagnetic Relaxation Enhancement for the Characterization of Transient Low-Population States of Biological Macromolecules and Their Complexes. Chemical Reviews 109, 4108–4139 (2009). 10.1021/cr900033p

44 Buratti, E. & Baralle, F. E. Characterization and functional implications of the RNA binding properties of nuclear factor TDP-43, a novel splicing regulator ofCFTR exon 9. Journal of Biological Chemistry 276, 36337–36343 (2001).

45 Mohanty, P., Rizuan, A., Kim, Y. C., Fawzi, N. L. & Mittal, J. A complex network of interdomain interactions underlies the conformational ensemble of monomeric TDP-43 and modulates its phase behavior. Protein Science 33, e4891 (2024). 10.1002/pro.4891

46 Arseni, D. et al. Heteromeric amyloid filaments of ANXA11 and TDP-43 in FTLD-TDP Type C. bioRxiv, 2024.2006.2025.600403 (2024). 10.1101/2024.06.25.600403

47 Alami, N. H. et al. Axonal transport of TDP-43 mRNA granules is impaired by ALS-causing mutations. Neuron 81, 536–543 (2014).

48 Fuentealba, R. A. et al. Interaction with polyglutamine aggregates reveals a Q/N-rich domain in TDP-43. Journal of Biological Chemistry 285, 26304–26314 (2010).

49 Skinner, S. P. et al. CcpNmr AnalysisAssign: a flexible platform for integrated NMR analysis. J Biomol NMR 66, 111–124 (2016). 10.1007/s10858-016-0060-y

50 Murthy, A. C. et al. Molecular interactions underlying liquid− liquid phase separation of the FUS low-complexity domain. Nature structural & molecular biology 26, 637–648 (2019).

51 Best, R. B., Zheng, W. & Mittal, J. Balanced Protein–Water Interactions Improve Properties of Disordered Proteins and Non-Specific Protein Association. Journal of Chemical Theory and Computation 10, 5113–5124 (2014). 10.1021/ct500569b

52 Abascal, J. L. F. & Vega, C. A general purpose model for the condensed phases of water: TIP4P/2005. The Journal of Chemical Physics 123, 234505 (2005). 10.1063/1.2121687

53 Luo, Y. & Roux, B. Simulation of osmotic pressure in concentrated aqueous salt solutions. The journal of physical chemistry letters 1, 183–189 (2010).

54 Páll, S. et al. Heterogeneous parallelization and acceleration of molecular dynamics simulations in GROMACS. The Journal of Chemical Physics 153, 134110 (2020). 10.1063/5.0018516

55 Evans, D. J. & Holian, B. L. The Nose–Hoover thermostat. The Journal of Chemical Physics 83, 4069–4074 (1985). 10.1063/1.449071

56 Berendsen, H. J., Postma, J. v., Van Gunsteren, W. F., DiNola, A. & Haak, J. R. Molecular dynamics with coupling to an external bath. The Journal of chemical physics 81, 3684–3690 (1984).

57 Zhang, Z., Liu, X., Yan, K., Tuckerman, M. E. & Liu, J. Unified Efficient Thermostat Scheme for the Canonical Ensemble with Holonomic or Isokinetic Constraints via Molecular Dynamics. The Journal of Physical Chemistry A 123, 6056–6079 (2019). 10.1021/acs.jpca.9b02771

58 Case, D. A. et al. *Amber* 2022. (University of California, San Francisco, 2022).

59 Darden, T., York, D. & Pedersen, L. Particle mesh Ewald: An N⋅ log (N) method for Ewald sums in large systems. The Journal of chemical physics 98, 10089–10092 (1993).

60 Ryckaert, J.-P., Ciccotti, G. & Berendsen, H. J. C. Numerical integration of the cartesian equations of motion of a system with constraints: molecular dynamics of n-alkanes. Journal of Computational Physics 23, 327–341 (1977). 10.1016/0021-9991(77)90098-5

61 Zheng, W. et al. Molecular Details of Protein Condensates Probed by Microsecond Long Atomistic Simulations. The Journal of Physical Chemistry B 124, 11671–11679 (2020). 10.1021/acs.jpcb.0c10489

62 Tribello, G. A., Bonomi, M., Branduardi, D., Camilloni, C. & Bussi, G. PLUMED 2: New feathers for an old bird. Computer physics communications 185, 604–613 (2014).

63 Courtney, N. J. et al. Insights into Molecular Diversity within the FET Family: Unraveling Phase Separation of the N-Terminal Low Complexity Domain from RNA-Binding Protein EWS. bioRxiv, 2023.2010.2027.564484 (2023). 10.1101/2023.10.27.564484

64 Pettersen, E. F. et al. UCSF Chimera—a visualization system for exploratory research and analysis. Journal of computational chemistry 25, 1605–1612 (2004).

65 Pettersen, E. F. et al. UCSF ChimeraX: Structure visualization for researchers, educators, and developers. Protein Science 30, 70–82 (2021).

