## Supplemental Information for "Structural details of helix-mediated TDP-43 C-terminal domain multimerization"

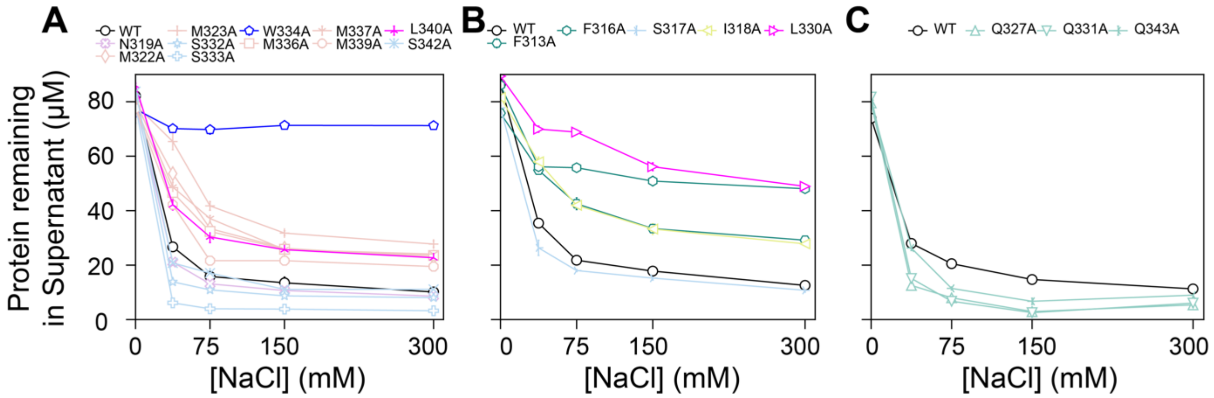


**Supporting Figure 1. Phase separation saturation concentrations of TDP-43 CTD and designed variants from alanine-scanning mutagenesis. A-C**. Quantification of phase separation assay showing the protein remaining in the supernatant after phase separation for WT CTD and its single alanine substitution variants at non-alanine positions within CR and its adjacent residues measured from 0 to 300 mM NaCl from the individual experiments that were performed together. Standard deviations of three replicates are represented as error bars.

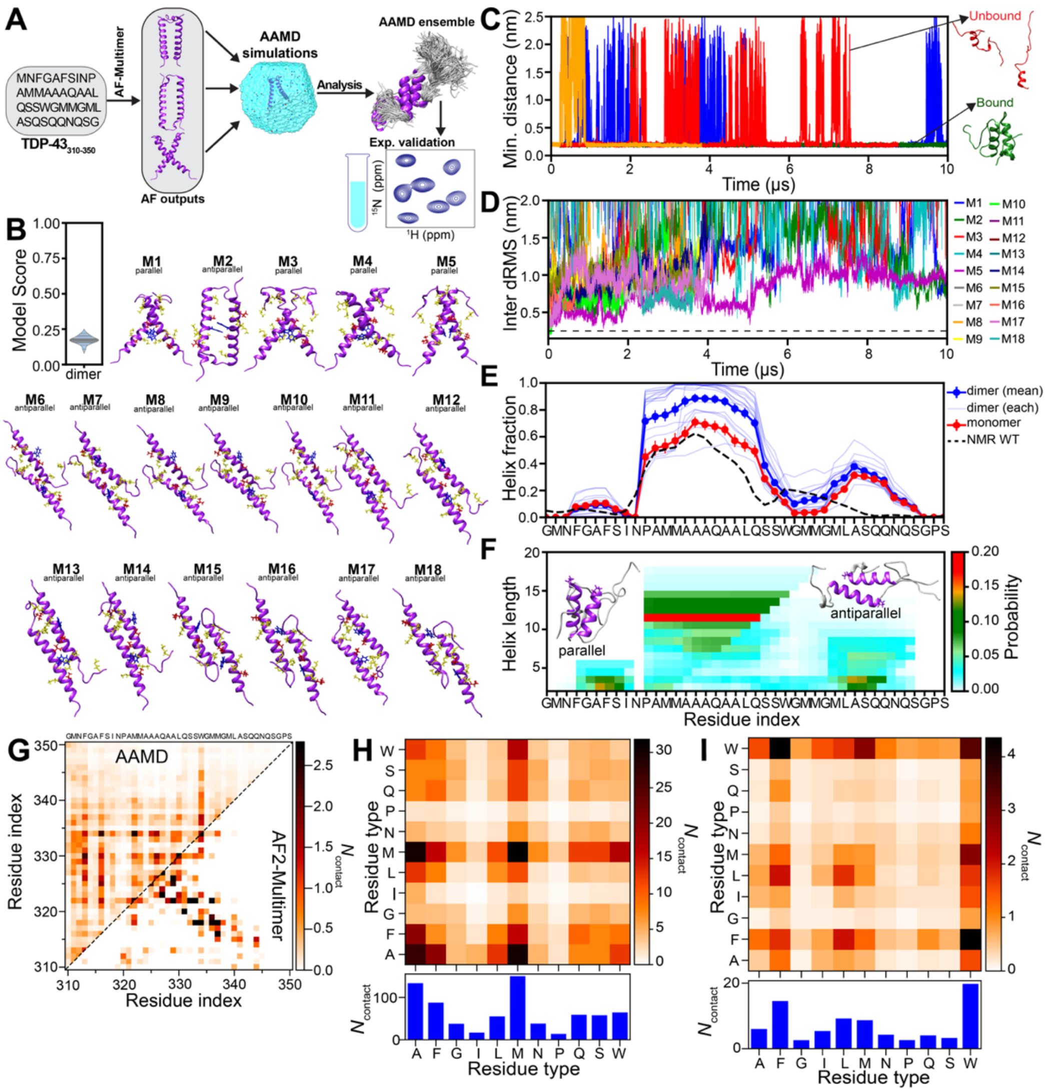


**Supporting Figure 2**. **Characterization of TDP-43$_{310–350}$ dimers from two-chain all-atom molecular dynamics (AAMD) simulations**. **A.** Schematics of simulation setup: Dimer structures of TDP-43$_{310–350}$ were predicted using AlphaFold-Multimer tool, then fed to run unbiased all-atom molecular dynamics simulations. These simulations were combined to form an ensemble (total runtime ~100 μs) of TDP-43$_{310–350}$ dimers for analysis and comparison to the experimental data. **B**. The model confidence score and initial structures of dimers predicted from AF2-Multimer used for two-chain atomistic simulations **C.** Minimum distance analysis between the TDP-43$_{310–350}$ dimers show their dynamic nature with dissociation and re-association of helices **D**. Inter-dRMS of all heavy atoms (excluding hydrogen atoms) for residues from 320 to 341 as a function of time with respect to initial conformations are computed from dimeric all-atom simulations. **E.** Per-residue helix fraction of TDP-43$_{310-350}$ chains, calculated from TDP-43$_{310-350}$ dimer (fractions from individual simulations shown as lighter color) and monomer simulations, compared with helix fractions from NMR-derived ensembles of TDP-43 CTD. **F**. Helix position and length map for TDP-43$_{310–350}$ show the highest probability of helices spanning from aa. 320 to 331. The representative snapshots corresponding to the most populated helix length show both parallel and antiparallel helix binding orientations. **G**. Pairwise intermolecular contact map as a function of residue index from two-chain AAMD simulations is compared to initial contacts in dimer structures from AF2-Multimer. **H-I.** Contact maps highlighting the average number of pairwise interactions between all residue types occurring in TDP-43$_{310–350}$ dimers from AAMD simulations shown in H (non-normalized) and I (normalized by the total count of each residue type occurring in the sequence), respectively.

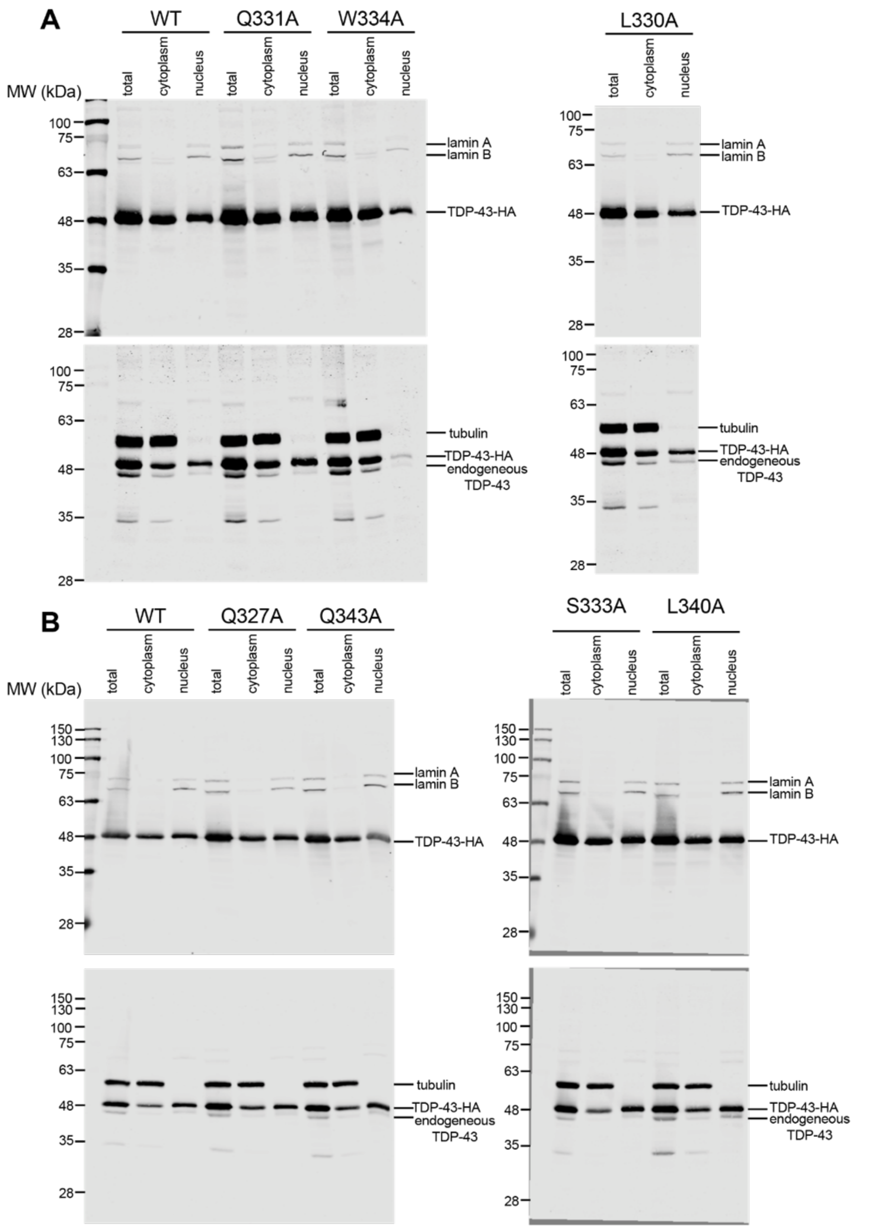


**Supporting Figure 3. TDP-43 CR helix assembly affects TDP-43 nuclear retention. A-B.** Representative immunoblot of total, cytoplasmic and nuclear fractions of HEK293$^{HA\text{-}TDP\text{-}43}$ cells from the individual experiments that were performed together. Lamin A/C and tubulin were used as nuclear and cytoplasmic controls, respectively.

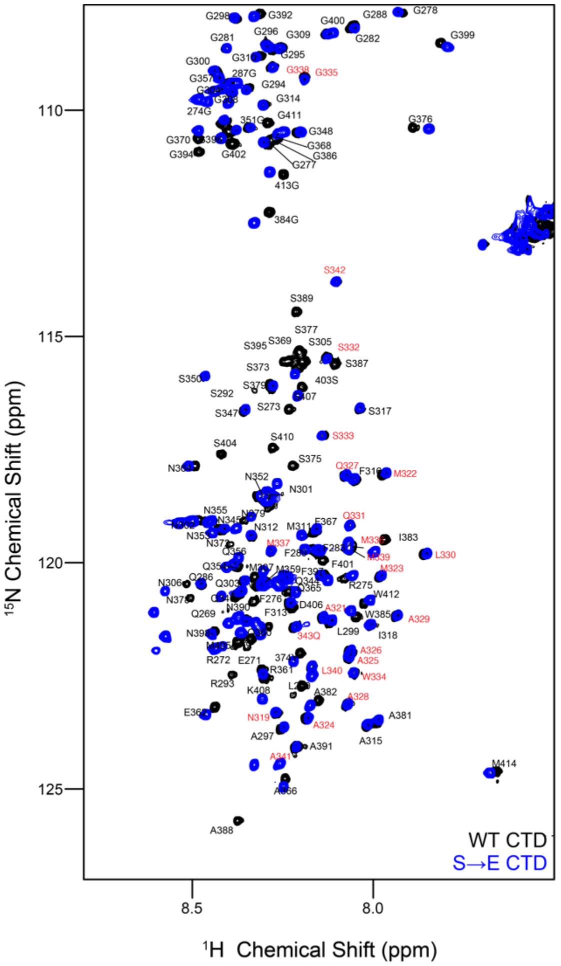


**Supporting Figure 4. Mutations in the flanking region dramatically enhance solubility. while preserving CR structure**. A comparison of the $^{1}$H-$^{15}$N HSQC of 20μM WT CTD (black) and 20 μM S→E CTD (blue) in 20 mM MES (pH 6.1) at 25ºC. The spectra are highly similar for the CR (red labels) where the resonances overlay nearly perfectly, suggesting no change in the structure of the CR is induced by the S→E substitutions.

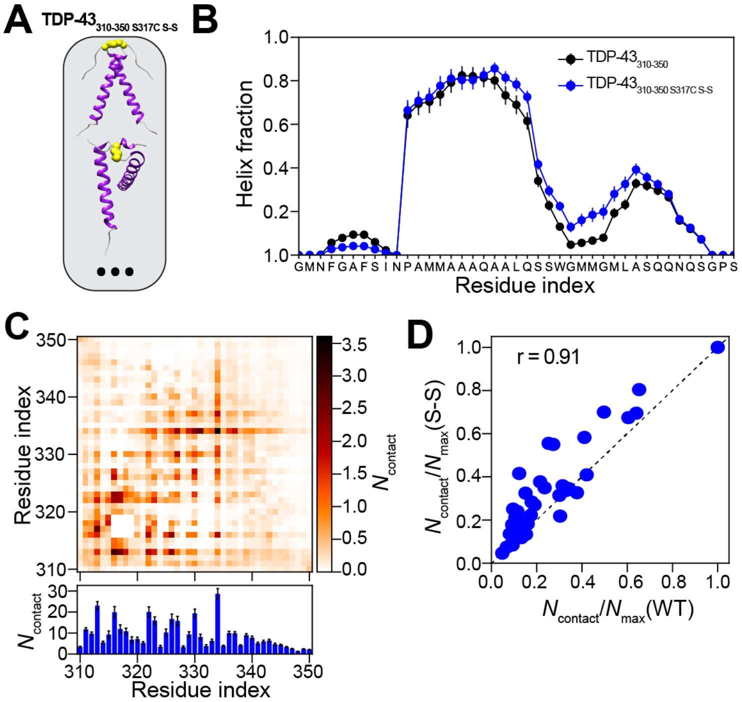


**Supporting Figure 5. Cross-linking near CR is expected to result in similar helicity and helix-helix contacts as in WT CTD dimers. A**. TDP-43$_{310-350}$ cross-linked dimers are modelled from AFv2.3 Multimer by adding a disulfide bond (S-S) at S317C. Then, 8 different starting conformations of the cross-linked dimers, TDP-43$_{310-350_S317C_S-S}$ are fed to run all-atom molecular dynamics simulations (3 independent runs for each model), resulting in a well-converged ensemble (total runtime ~48 µs). **B**. Per–residue helix fraction of TDP-43$_{310-350}$ chains calculated from TDP-43$_{310-350}$ dimer (WT) and TDP-43$_{310-350_S317C_S-S}$ cross-linked dimer simulations. TDP-43 cross-linked dimer shows helix content similar to those observed in the wild-type. **C**. A two-dimensional pairwise intermolecular contact map as function of residue index (top) and total number of intermolecular contacts per residue position ($N_{contact}$) (bottom) is computed from of the simulated cross-linked dimer ensemble. **D**. Per-residue intermolecular contacts (normalized by maximum contact in each case) calculated from TDP-43$_{310-350}$ (WT) and cross-linked (S-S) dimer simulations show a strong correlation (Pearson correlation coefficient, r = 0.91).

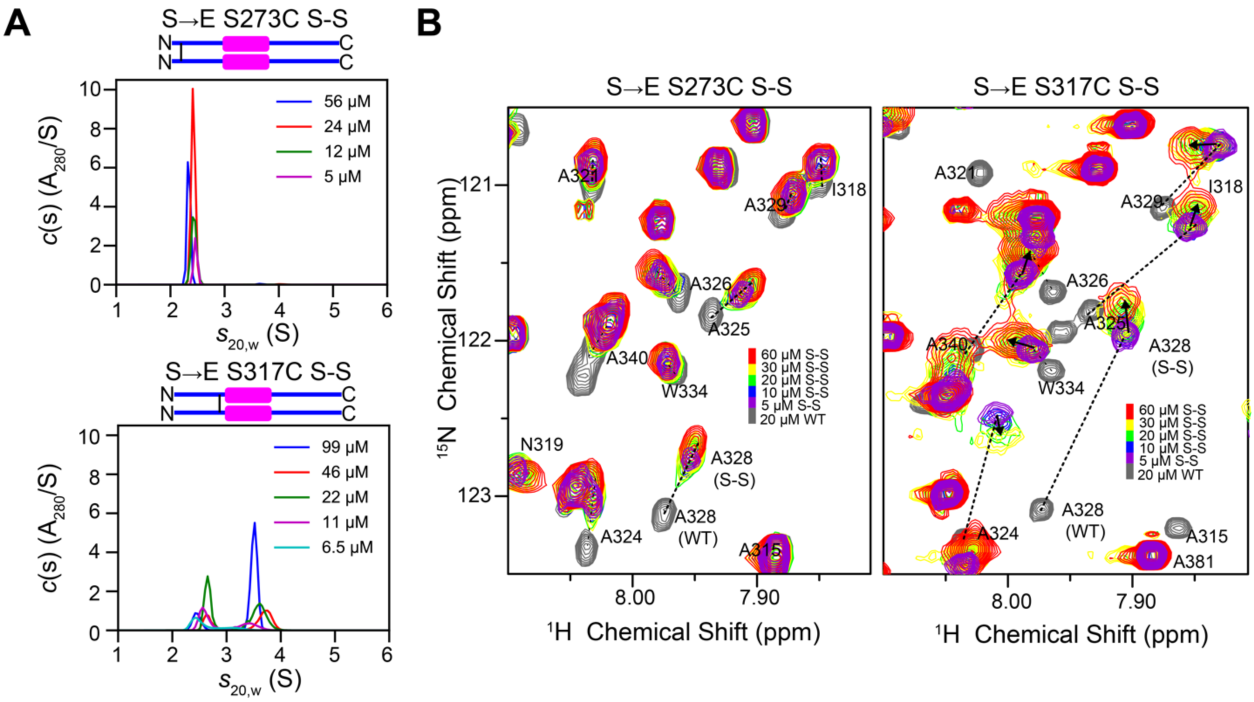


**Supporting Figure 6**. **TDP-43 CTD forms higher-order oligomers larger than dimer A**. Sedimentation velocity analytical ultracentrifugation (SV-AUC) of cysteine cross-linked variants (S→E S273C and S→E S317C) with increasing protein concentrations, at rotor speeds of 5000 rpm, 25ºC. SV-AUC data were analyzed as a continuous *c*(s) distribution of sedimenting species. Sedimentation coefficients were corrected to standard conditions at 20 °C in water, $s_{20,w}$. SV-AUC suggests the presence of species larger than dimer (~2.4 S - 31.5 kDa). **B**. NMR chemical shift analysis of cysteine cross-linked variants (S→E S273C and S→E S317C) as a function of increasing concentration shows change in chemical environment compared to monomeric WT CTD (20 μM), suggesting the formation of higher order oligomers.

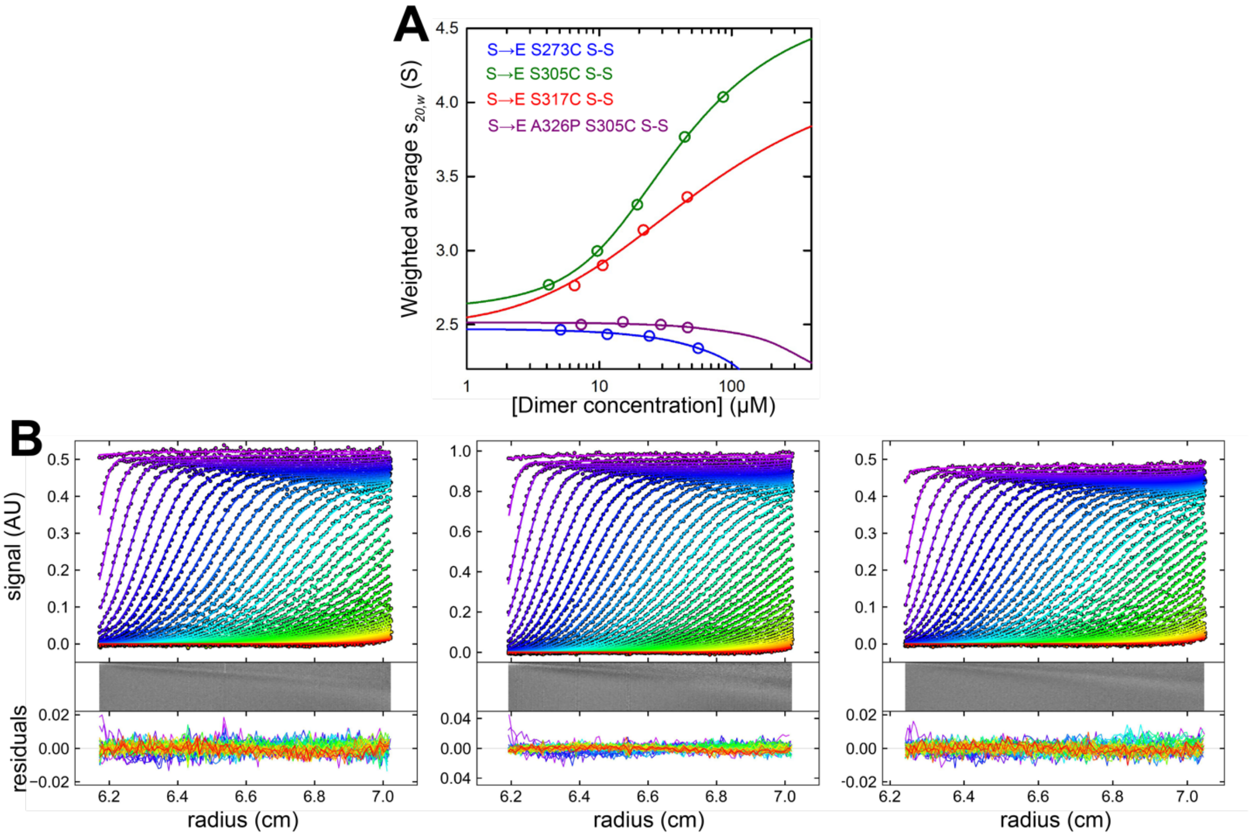


**Supporting Figure 7**. **A**. Weighted average sedimentation coefficient isotherms as a function of protein concentration based on the sedimentation velocity c(s) distributions. S→E S273C (blue) and S→E S305C A326P (purple) presented as dimers at all concentrations studied and a linear fit was used to describe the concentration dependence of the sedimentation coefficient. S→E S317C (red) were modeled in terms of a dimer – tetramer reversible self-association to obtain a dimer – tetramer $K_d$ of 46 μM (in dimer concentration units). S→E S305C (green) were modeled in terms of a dimer – tetramer – octamer reversible self-association to obtain a dimer – tetramer $K_d$ of 58 μM and a tetramer – octamer $K_d$ of 4.7 μM, indicative of cooperative self-assembly. **B.** Absorbance sedimentation data collected for S→E S317C at 280 nm and (left) 47 μM (3 mm pathlength cell), (center) 22 μM (12 mm pathlength cell), and (right) 11 μM (12 mm pathlength cell) were analyzed globally in terms of a dimer – tetramer self-association using Lamm equation modeling, along with trace amounts of an aggregate. The analysis, carried out in SEDPHAT, returns a dimer – tetramer $K_d$ of 46 μM (68% confidence interval of 42 – 51 μM). Data were plotted in GUSSI and for clarity only every third scan and every third experimental data point are shown. Best-fits are represented by a solid line through the experimental points. A bitmap representation of the residuals, together with the combined residuals, are shown below each plot.

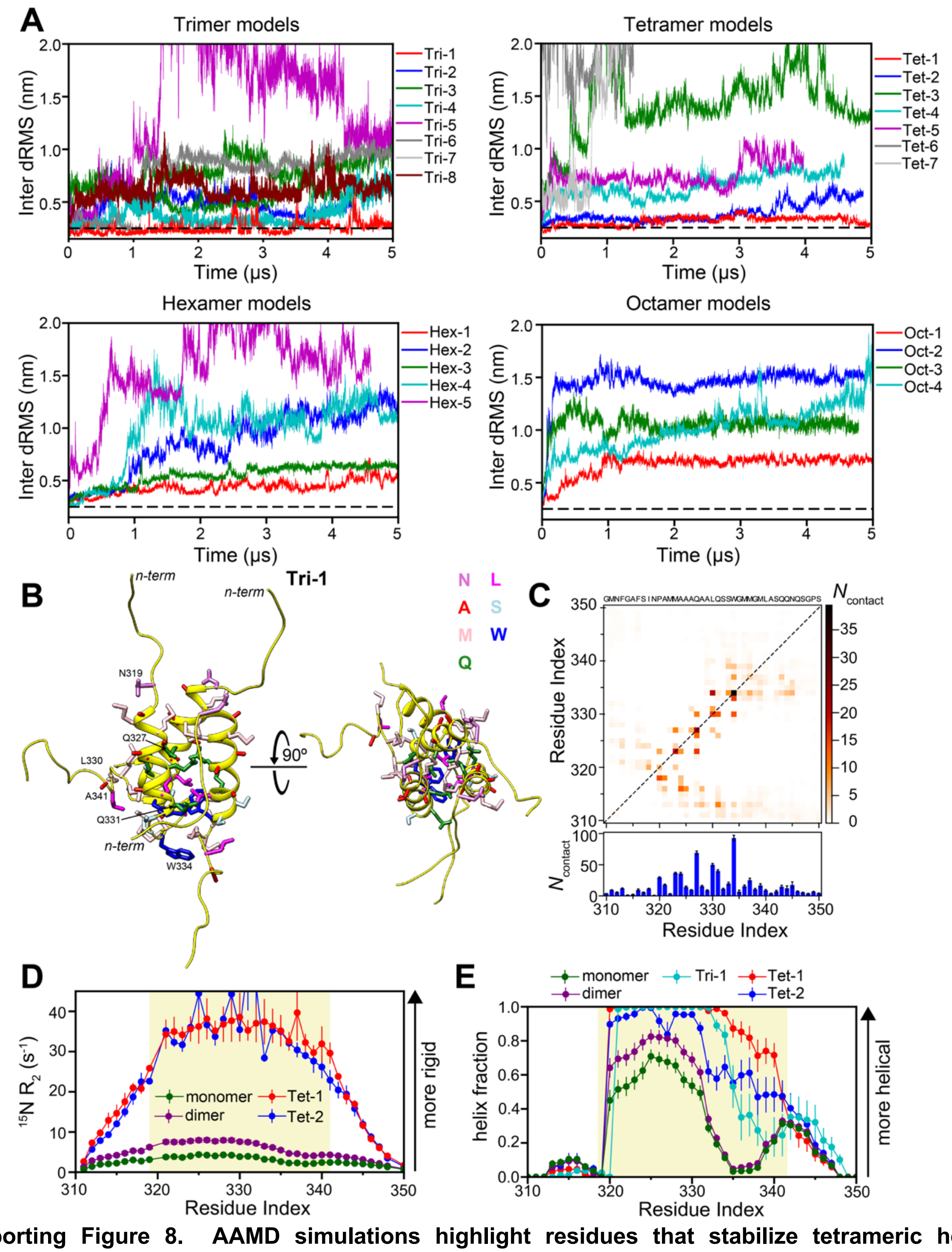

**Supporting Figure 8. AAMD simulations highlight residues that stabilize tetrameric helical assembly of TDP-43 CTD. A.** Inter-dRMS analysis of oligomeric models computed from the AAMD simulations (5 μs each) of AlphaFold-Multimer predictions. Inter-dRMS of all heavy atoms (excluding hydrogen atoms) for residues from 320 to 341 as a function of time with respect to initial conformations are shown. Intermolecular contacts are lost compared to initial PDB contacts in both hexamer and octamers models. One structure (Tri-1) is found to be stable among trimer models, whereas Tet-1 and Tet-2 exhibit greater stability compared to other tetramer models. **B**. The most representative structure of Tri-1 from the last microsecond of AAMD simulations (single run, 5 μs) are shown. Helices are shown as yellow-colored ribbons. Side chainx (excluding hydrogen atoms) of CR residues are shown as sticks, with their colors indicating different residue types as illustrated in the right corner. **C**. (top) Pairwise intermolecular contact map of Tri-1 from AAMD simulations (single run, 5 μs). (bottom) Total number of contacts per residue position ($N_{contact}$) derived through summation of all pairwise contacts along y-axis based on two dimensional pairwise intermolecular contact maps. **D.** $^{15}N$ $R_2$ of Tet-1 and Tet-2 from AAMD simulations (single run, 5 μs) shows slower motions compared to monomeric and dimeric AAMD ensembles, consistent with experiments. **E.** Per-residue α-helical fraction from Tet-1 and Tet-2 AAMD simulations (single run, 5 μs) show increase in helicity in the main helical region (aa: 320-331) and further helix extension in the adjacent 332–343 region with helix-helix assembly compared to monomeric and dimeric AAMD ensembles and Tri-1 AAMD simulation (single run, 5 μs).

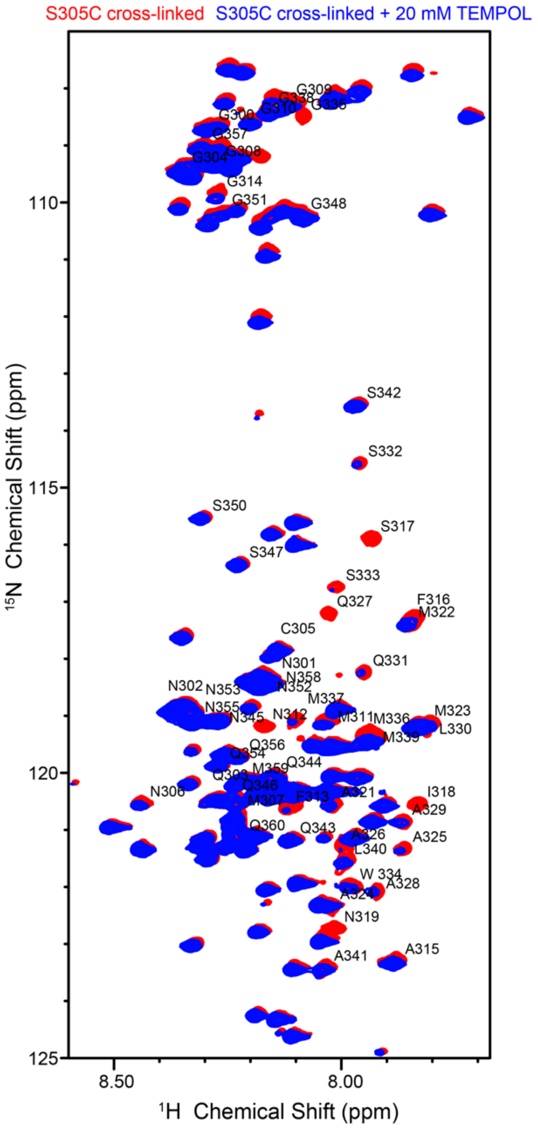


**Supporting Figure 9. Solvent PRE analysis of TDP-43 S$\rightarrow$E S305C cross-linked dimer variant**. $^1H$-$^{15}N$ HSQC of 30 μM S$\rightarrow$E S305C cross-linked dimer without (red) and with 20 mM TEMPOL (blue) at 47ºC, 850 MHz

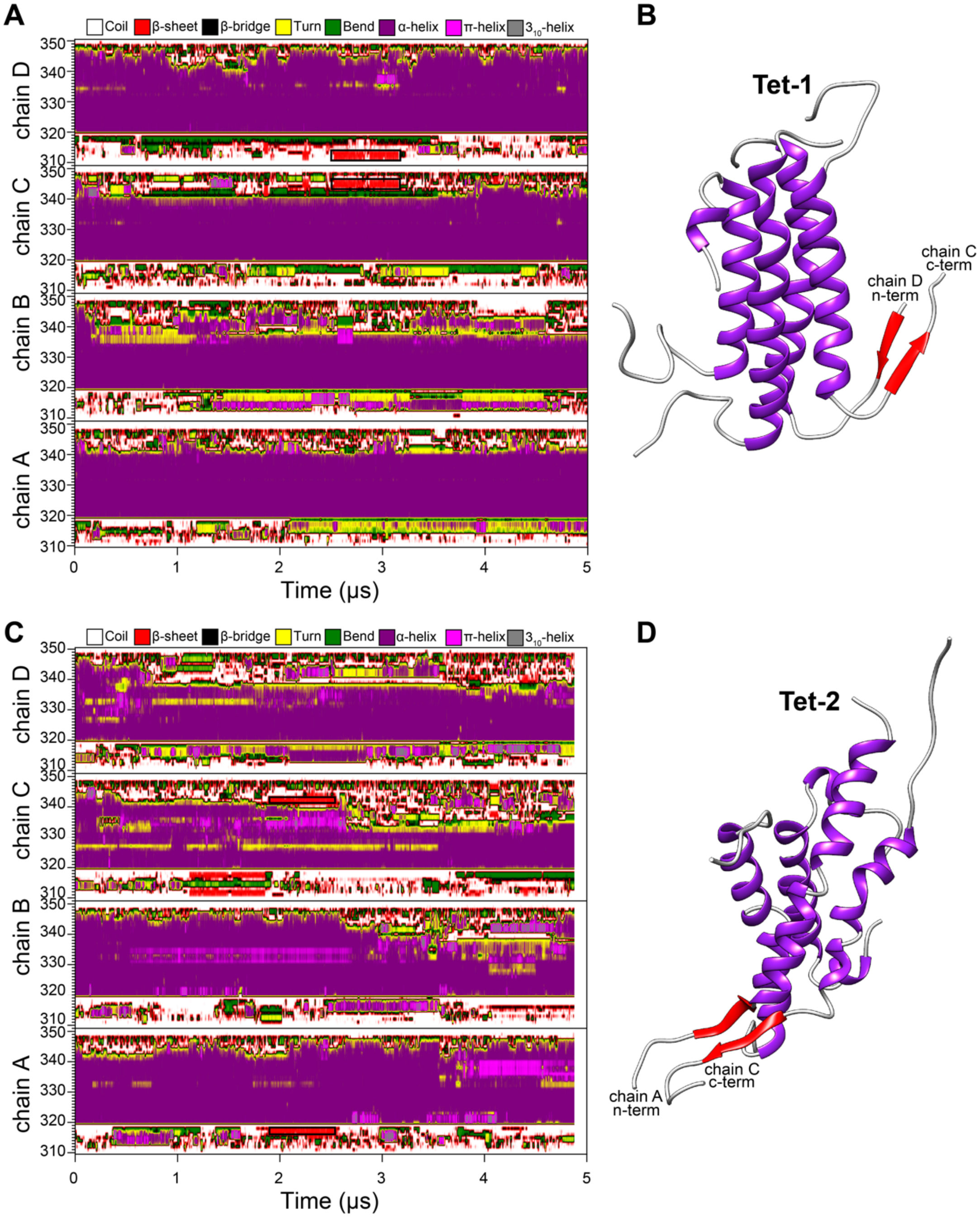


**Supporting Figure 10. Secondary structure analysis of TDP-43 CR tetrameric assemblies from AAMD simulations**. Secondary structure change as a function of time in Tet-1 AAMD simulation trajectory (single run, 5 μs) shows the formation of intermolecular β-sheet-structure (highlighted in black boxes). **B**. The snapshot from Tet-1 AAMD simulation that shows the formation of transient, antiparallel β-sheets (red strands) between residues $^{310}$GMNF$^{313}$ of chain C and residues $^{344}$QNQ$^{346}$ of chain D. **C**. Secondary structure change as a function of time in Tet-2 AAMD simulation trajectory (single run, 5 μs) shows the formation of intermolecular β-sheet-structure (highlighted in black boxes). **D**. The snapshot from Tet-2 AAMD simulation that shows the formation of transient, antiparallel β-sheets (red strands) between residues $^{315}$AFSI$^{318}$ of chain A and residues $^{341}$ASQQN$^{345}$ of chain C.

**Table S1**. Serine positions in TDP-43 CTD simultaneously changed to glutamate in S→E variant series used in NMR experiments.

| Mutant | Residues |
|---|---|
| S→E | S292, S305, S369, S377, S379, S387, S389, S393, S395, S403, S404, S409, S410 |

**Table S2:** Oligonucleotides used in these studies.

| DNA | Sequence |
|---|---|
| TDP43_Q327A Fw | 5'-GCCATGATGGCTGCCGCCGCTGCAGCACTACAGAGCAGTTGG-3' |
| TDP43_Q327A Rv | 5'-CCAACTGCTCTGTAGTGCTGCAGCGGCGGCAGCCATCATGGC-3' |
| TDP43_Q331A Fw | 5'-GCCGCCCAGGCAGCACTAGCTAGCAGTTGGGGTATGATG-3' |
| TDP43_Q331A Rv | 5'-CATCATACCCCAACTGCTAGCTAGTGCTGCCTGGGCGGC-3' |
| TDP43_Q343A Fw | 5'-ATGGGCATGTTAGCCAGCGCTCAGAACCAGTCAGGCCCATCG-3' |
| TDP43_Q343A Rv | 5'-CGATGGGCCTGACTGGTTCTGAGCGCTGGCTAACATGCCCAT-3' |
| TDP43_L330A Fw | 5'-GCTGCCGCCCAGGCAGCAGCTCAGAGCAGTTGGGGTATG-3' |
| TDP43_L330A Rv | 5'-CATACCCCAACTGCTCTGAGCTGCTGCCTGGGCGGCAGC-3' |
| TDP43_W334A Fw | 5'-GCAGCACTACAGAGCAGTGCTGGTATGATGGGCATGTTAG-3' |
| TDP43_W334A Rv | 5'-CTAACATGCCCATCATACCAGCACTGCTCTGTAGTGCTGC-3' |
| TDP43_S333A Fw | 5'-CAGGCAGCACTACAGAGCGCTTGGGGTATGATGGGCATG-3' |
| TDP43_S333A Rv | 5'-CATGCCCATCATACCCCAAGCGCTCTGTAGTGCTGCCTG-3' |
| TDP43_L340A Fw | 5'-TGGGGTATGATGGGCATGGCTGCCAGCCAGCAGAACCAG-3' |
| TDP43_L340A Rv | 5'-CTGGTTCTGCTGGCTGGCAGCCATGCCCATCATACCCCA-3' |
